# Community Challenge towards Consensus on Characterization of Biological Tissue: C^4^Bio’s First Findings

**DOI:** 10.1101/2025.07.30.663632

**Authors:** Nele Famaey, Heleen Fehervary, Yoann Lafon, Ali Akyildiz, Silke Dreesen, Karine Bruyère-Garnier, Jean-Marc Allain, Marta Alloisio, Alejandro Aparici-Gil, Chiara Catalano, Fanette Chassagne, Snehal Chokhandre, Kimberly Crevits, Hanneke Crielaard, Eoghan Cunnane, Connor Cunnane, Karen De Leener, Amisha Desai, Rob Driessen, Ahmet Erdemir, Mona Eskandari, Sam Evans, Christian Gasser, Marc Gebhardt, Birgit Glasmacher, Gerhard A. Holzapfel, Mikel Isasi, Louise Jennings, Sascha Kurz, Sara Leal-Marin, Pauline Lecomte, Annie Morch, John Mulvihill, Fulufhelo Nemavhola, Thanyani Pandelani, Salvatore Pasta, Estefania Peña, Baptiste Pierrat, Heidi-Lynn Ploeg, Stanislav Polzer, Manuel Rausch, David Schwarz, Hazel Screen, Selda Sherifova, Gerhard Sommer, Shengzhang Wang, Darragh Walsh, Deepesh Yadav, Thierry Marchal, Liesbet Geris

## Abstract

This study investigates methodological variability across various expert laboratories worldwide, with regards to characterizing the mechanical properties of biological tissues. Two testing rounds were conducted on the specific use case of uniaxial tensile testing of porcine aorta. In the first round, 24 labs were invited to apply their established methods to assess inter-laboratory variability. This revealed significant methodological diversity and associated variability in the stress-stretch results, underscoring the necessity for a standardized approach.

In the second round, a consensus protocol was collaboratively developed and adopted by 19 labs in an attempt to minimize variability. This involved standardized sample preparation and uniformity in testing protocol, including the use of a common cutting and thickness measurement tool. Despite protocol harmonization, significant variability persisted across labs, which could not be solely attributed to inherent biological differences in tissue samples.

These results illustrate the challenges in unifying testing methods across different research settings, underlining the necessity for further refinement of testing practices. Enhancing consistency in biomechanical experiments is pivotal when comparing results across studies, as well as when using the resulting material properties for in silico simulations in medical research.

## 1. Introduction

Characterization of the mechanical properties of biological tissue serves many purposes. Firstly, it can help to improve our fundamental understanding of physiological function, due to a strong structure-function relationship for many organs and tissue systems. Secondly and as a consequence of the former, ageing as well as various diseases are known to alter the mechanical properties of the involved tissues, such that quantification of these alterations can aid diagnosis and monitoring. Thirdly, in the field of tissue engineering and regenerative medicine, it is imperative that the engineered tissues replicate the mechanical behaviour of native tissues to ensure proper functioning. Likewise mechanical data informs the design of implants, prosthetics, and surgical tools to ensure compatibility with tissue mechanics and minimize adverse reactions. Indirectly, mechanical properties can be used to inform constitutive models which can in turn be used in computational models, for various applications in the field of *in silico* medicine (Motiwale & Sacks, 2025). However, although the scientific literature therefore abounds with articles experimentally characterizing the mechanical properties of biological tissues, there are still no widely recognized testing standards for these experiments. This shortcoming has consequences for the interpretability of the results, especially when comparing across studies. Moreover, when this experimental data is used to derive material properties used as input parameters to *in silico* models, the associated error and uncertainty propagates into these simulations.

Simulation-driven medical device development and in silico trials are gaining importance (Pappalardo et al., 2022; Viceconti & Emili, 2024). Clearly, the quality of these simulations is of utmost importance if *in silico* medicine is to take its rightful place in medical research and development. Consequently and analogously to the well-known concepts of ‘good medical practice’ and ‘good laboratory practice’ (Robertson & Williams, 2009; Stevens, 2003), there is a growing emphasis on the development of guidelines for ‘good simulation practice’. In 2018, the American Society of Mechanical Engineers introduced a standard known as ‘ASME V&V 40 - Assessing Credibility of Computational Modeling through Verification and Validation: Application to Medical Devices’ (ASME, 2018), and in 2023, the FDA issued a guidance document entitled ‘Assessing the Credibility of Computational Modeling and Simulation in Medical Device Submissions’ (FDA, 2023). For example, a reporting checklist for verification and validation of finite element analysis was made more specifically for orthopaedic and trauma biomechanics (Oefner et al., 2021).

Such guidelines span various aspects related to model development, verification, calibration and validation, and focus their attention on the quantification of the uncertainty that accumulates in each of these aspects. Apart from modeling assumptions and discretization errors, it is intuitive that model input data, whether used to calibrate model parameters or validate the model outcomes, are an important source of uncertainty. Indeed, credible numerical simulations require input parameters and underlying acquisition methods that are both traceable and reliable (ASME, 2018). However, where numerical analysts are usually well-equipped to test the validity of their modeling assumptions or to minimize discretization errors, not every numerical analyst has the expertise and/or facilities to assess the quality and uncertainty of their model input data, especially when it involves data related to biological tissue properties.

Ideally, the required material properties and their uncertainty should come from shared, trusted databases. However, various challenges are faced when attempting to create such a database. Firstly, depending on the context of use of a simulation (Viceconti et al., 2021), different types of material properties might be required (mechanical, thermal, electrical, etc.). Even when focusing on mechanical properties alone, a simulation might require properties related to a specific material constitutive law and its characteristics, such as the tensile elasticity, viscoelasticity, anisotropy, compressibility, nonlinearity, ultimate tensile strength, pre-stress state etc. Each of these properties can, in turn, be acquired by various test setups and methods, such as uniaxial or biaxial tensile tests, compression tests, shear tests, indentation tests, dynamic mechanical analysis, and creep tests. The selection of the appropriate testing method depends on the tissue type, its anatomical location, and the desired application/objective to which the simulation responds. Moreover, apart from the actual testing method, the sample collection, preservation and preparation method will also influence the resulting properties, even if such effects are still under debate (Blaker et al., 2024; Oswald et al., 2017; Chow & Zhang, 2011; Stemper et al., 2007).

The second challenge relates to the population- and subject-related variability. Indeed, material properties of biological tissues exhibit significant variability due to physiological and demographic factors. Sex-related variations and ethnic diversity contribute to changes in tissue composition and structure, leading to differences in material properties for a given tissue type (Smoljkić et al., 2023; Åstrand et al., 2011). Age-related changes, such as decreased elasticity and increased brittleness in tissues like skin and bone, or other pathological conditions, induce further variability (Kirilova-Doneva & Pashkouleva, 2022; Mirzaali et al., 2016). Even within a subject, properties will vary depending on the location within the body (Krueger et al., 2011). Ideally, a material properties database would encompass this broad representation, which would require the collection of an extremely large range of experimental data. Moreover, in certain research communities, there appears to be an unsubstantiated trust in the ‘known values’ taken from the literature. As Hammer & Klima (2019) also noted in their review, the implementation of further region-specific studies is partially inhibited, as reliance is placed on a supposedly existing broad database, instead of carrying out new studies adapted to the research question. Using numerical studies of the biomechanics of the sacro-iliac joint as an example, they point out the frequent nested literature references and the often accompanying decrease in relevance of the basic source for the intended loading scenario.

With that in mind, perhaps the most important challenge related to building up such a database is related to methodological variability. Indeed, even when comparing literature results of the same tissue type of a similar animal strain or patient group, tested according to - reportedly - the same method, results vary widely. As mentioned, there are currently no established standards for measuring the material properties of biological tissue, nor is there a unified approach for sample preparation and storage, or for reporting and assessing the obtained results. Some groups have proposed guidelines for specific aspects or applications, e.g. Kurz et al. (2023) have recently suggested a standardized approach for characterization of the human lumbopelvic system, Wale et al. (2021) have investigated sample preparation and optimal sample shape to induce reproducible failure for tensile testing of musculoskeletal soft tissues, Scholze et al. (2020) have proposed a clamping system for simple and reproducible sample clamping. Though valuable, these studies are limited to a single research group, and it remains to be seen if and how they will be adopted by the scientific community. Lin et al. (2024) have recently performed a systematic review of uniaxial tensile testing of human soft tissues across research groups. They found large variations in sample shape and especially underreporting of various aspects of the protocol, including the clamping mechanism. Similarly, Fehervary (2018) investigated variations in planar biaxial testing of arterial tissue, both in experimental setup and data processing, and formulated guidelines for testing and reporting thereof. Although this lack of standardization and high variability is commonly known, to the best of our knowledge, a quantitative analysis of the current degree of methodological variability has not yet been attempted. It is therefore an essential first step towards standardization, ultimately leading to reliable material properties, including their uncertainty, which can be used in computational analyses for *in silico* medicine.

In 2019, the Virtual Physiological Human institute together with the Avicenna Alliance launched a task force on tissue characterization, out of which the C^4^Bio (C^4^Bio, 2025) initiative was born, an acronym for ‘Community Challenge towards Consensus on Characterization of Biological Tissue’. Considering the wide range of properties, tissue types and testing methods, the decision was made to direct the initial focus to ‘simple’ uniaxial tensile testing (i.e. the method most commonly found in literature) of porcine aorta. This paper first describes the methods employed to compare the results obtained by 24 expert laboratories from around the globe who characterized biological tissues and synthetic samples during two test rounds. In a first testing round, laboratories were instructed to use their own established method, to allow an unbiased evaluation of existing methodologies. In a second testing round, participants worked together to create a ‘consensus methodology’ and were then instructed to perform their tests accordingly.

## 2. Materials & methods

### 2.1. Participant recruitment & overall methodology

The initiative was announced in the fall of 2020 through various communication channels. All research groups with experience in material characterization of biological tissue were invited to join, provided the following prerequisites:

- the availability of infrastructure suitable for uniaxial tensile testing of samples within a length range of 10-60 mm and a linearized Young’s modulus range of 0-5 MPa,
- the clearance to test biological tissue in the laboratory,
- a demonstrated experience in mechanical experiments on biological tissue through scientific references,
- the willingness to publicly share the experimental results.

In round 1, 24 laboratories from all over the world participated (see Figure S1 in the supplementary material, section Appendix A, for a map), and 19 of them further participated in round 2. The participants that were no longer able to participate indicated that this was due to a lack of budget and/or person power.

For both testing rounds, all samples were procured and prepared centrally and shipped as described in Section 2.2. In round 1, 24 laboratories performed mechanical testing and analysis according to their preferred methodology, with only a limited number of instructions given (see Section 2.3). After testing, raw and processed data were collected and analysed centrally according to the methods described in Section 2.4. When providing the raw and the processed data, participants were also asked to fill in a survey, querying on various aspects of the testing methodology they used. Following the central analysis, the observations were discussed with the participants over multiple online meetings, and a consensus protocol was established and used in round 2. The full protocol as shared with the participants can be found in the supplementary material (section Appendix A).

### 2.2. Sample preparation & shipping

#### 2.2.1. Biological tissue

In both testing rounds, 2 sets of a proximal and a distal segment of one porcine descending aorta was prepared per participant at FIBEr, the KU Leuven core facility for Biomechanical Experimentation. Each round, aortas were collected from 70 animals from a local slaughterhouse, and brought to FIBEr in physiological medium (saline solution) at 4^◦^C. Care was taken to minimize biological variability, by harvesting material as much as possible from animals of the same strain, average weight and age, and distributing these randomly over the participants. From each aorta a distal and proximal segment of approximately 2 cm in diameter and 5 cm in length was cut, with a small proximal anterior incision, to allow tracking of the orientation of the sample (see Figure S2 in the supplementary material, section Appendix A). Samples were placed in individual containers with physiological medium, while keeping track of the animal and the location (proximal vs distal). All samples were transferred to a -80^◦^C freezer inside the containers within 8 hours of excision from the animal, and stored in FIBEr until shipment. The samples were shipped to each of the participants on dry ice in styrofoam boxes. Participants were instructed to keep the aortic tissue in frozen conditions (i.e. at -80^◦^C) until they were ready for testing, or to start the thawing and testing process immediately if no freezing capability was available in the laboratory. Upon receipt of the package, the participants verified that a sufficient amount of dry ice was still present in the box to ensure proper sample preservation. For all samples, the duration of frozen storage was recorded.

#### 2.2.2. Synthetic samples

In round 1, two sheets of synthetic elastomer material of approximately 4.5x9cm were shipped to the participating research group along with the biological tissue, in casu ‘Leartiker-K73’ and ‘Leartiker-K51’, provided by Leartiker s.a. (Spain).

In round 2, to use a material with more biologically relevant properties as compared to round 1, samples were 3D printed by Materialise NV (Belgium) using the company’s proprietary HeartPrintFlexPlus (HPF+) material (Schickel et al., 2019) in the dogbone shape as agreed upon in the consensus protocol (see section 2.3.2). In both rounds, the synthetic material was kept and shipped in dry conditions at ambient temperature.

### 2.3. Mechanical testing

#### 2.3.1. Round 1

In the first testing round, only minimal instructions were given to participants. This was done on purpose to allow for an objective comparison of the participating laboratories preferred methodologies. It was requested that all tests were executed by the same person, to avoid any inter-user variability. Also, the same equipment and methodology had to be used for both the aortic and the synthetic samples. The only exception in the process was that the synthetic material should always be tested in dry conditions.

The evening before the testing day, the participating groups placed the aortic specimen in a fridge (4°C) to thaw overnight. This allowed a thawing time of around 16 h, whereby the exact time per specimen was recorded. Next, the specimens were divided into test samples. From each tubular segment, at least 3 uniaxial test samples were prepared, with geometry and cutting method varying per participant. Proximal and distal segments were used to prepare circumferentially and longitudinally oriented samples, respectively. Participants were instructed to take a picture or make a sketch to indicate the location of each of sample with respect to the original specimen.

For the synthetic specimens, at least 6 samples were cut out of each provided sheet, in a direction of choice since the material is considered to be isotropic. Figure S3 in the supplementary material (section Appendix A) gives an overview of all samples (biological and synthetic) tested per participant.

For each (biological and synthetic) sample, the undeformed sample thickness, undeformed sample width in the neck region, and undeformed length of the neck region were measured according to each participant’s methodology.Finally, participants also selected individually the appropriate loading protocol and data processing methodology to determine the elastic stiffness moduli at various strain levels and the ultimate tensile strength.

#### 2.3.2. Round 2

In round 2, the same thawing instructions were given as in round 1. After thawing, test samples were excised from the aortic segments and at least three circumferentially oriented samples were prepared from the proximal segments and three longitudinally oriented samples from the distal segments. This time, the samples were cut into a dogbone shape using a cutting tool that was shipped to the participating research group along with the tissue. The dogbone shape and cutting tool are shown in Figure 1a and b and more details are available in 2.2. The time required for each phase in the preparation and testing protocol was logged for each sample.

**Figure 1:**
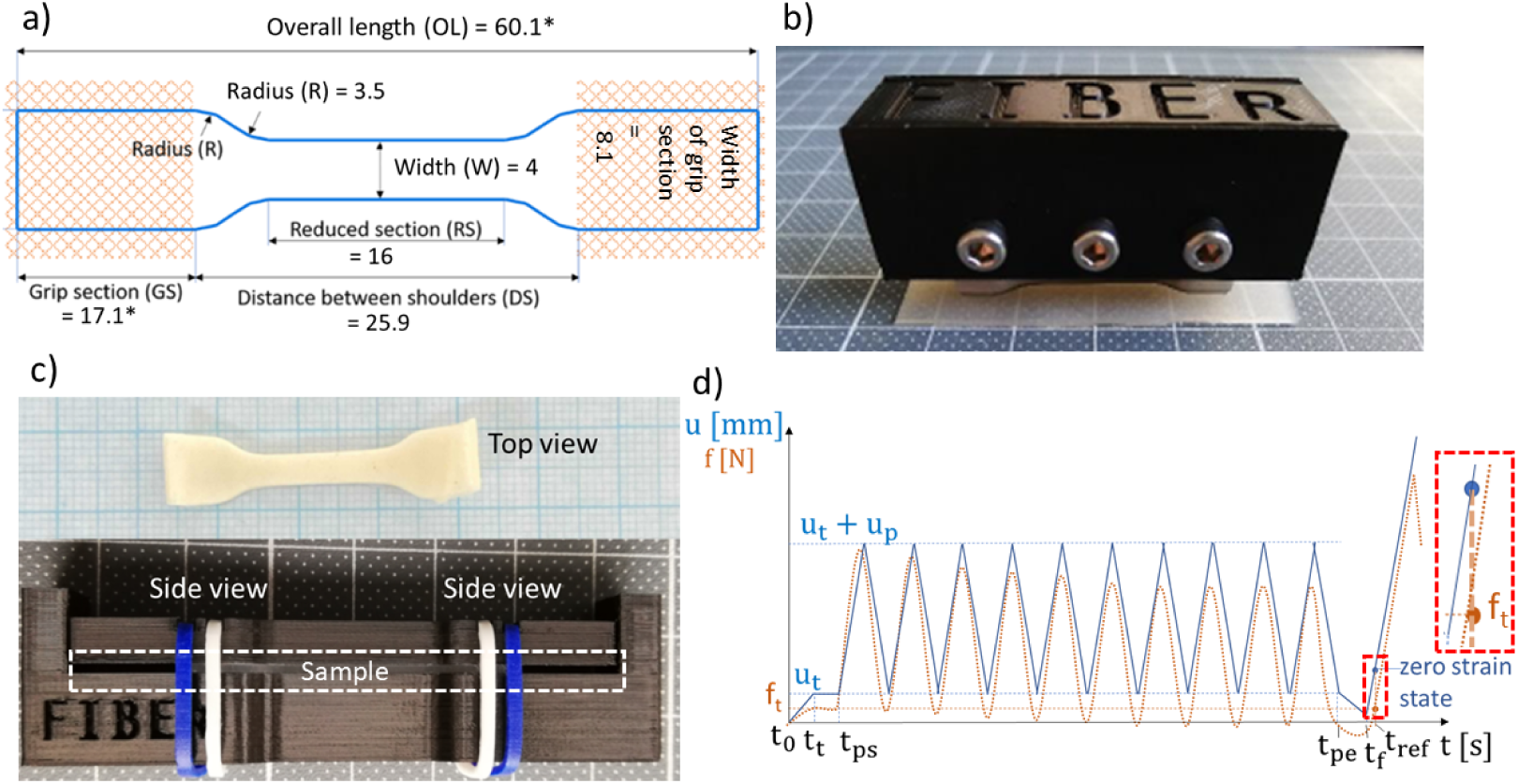
In round 2, a) samples were cut into a dogbone shape as agreed upon by all participants. *These dimensions can be increased if more gripping surface is required. All dimensions are in millimeter. b) A cutting tool was provided to ensure consistency in the shape. c) Calibrated pictures were taken during sample preparation for dimension measurements. Top: aortic sample on millimeter paper. Bottom: synthetic sample in a custom-made thickness measurement tool, tightened with elastic bands. d) Loading protocol for round 2, showing actuator displacement *u* and expected reaction force *f* as a function of time *t*. Time *t_t_* corresponds to the time at which the threshold force of *f_t_* is reached for the first time. Times *t_ps_* and *t_pe_* correspond to the start and end of the preconditioning cycles, during which a preload amplitude *u_p_* is applied for 10 cycles. Time *t_f_* corresponds to the start of the final upward ramp and *t_ref_* corresponds to the assumed zero strain state.

Participants were required to measure the undeformed width in the neck region and overall length of each sample using calibrated images taken with a high-resolution camera on millimeter paper. For the thickness measurement, the sample was inserted into a tool that was provided to the participants, shown in Figure 1c. This tool gently squeezes the sample with the help of identical elastic bands between two plates with known dimensions, such that the sample thickness can be derived from a side view image focussed on the tissue surface.

Participants were advised to use digital image correlation (DIC) to track sample deformation during the test, which required application of a speckle pattern on the sample’s intimal surface, by sprinkling graphite powder on a piece of paper and subsequently placing the intimal side of the sample onto the paper. Alternatively, if no DIC system was available, participants used a marker tracking technique, placing four markers at ∼6 mm intervals in the central area of the sample.

Similar to round 1, the participants were asked that all samples were tested using the same equipment by a single operator. After sample acclimatisation in saline solution at 37°C for approximately 10 minutes, the load cell was zeroed. For a horizontal set-up, this was done before mounting the sample, whereas for a vertical set-up, this was done after the sample was fixed in the upper clamp. During testing, the biological samples were immersed in a saline bath at 37°C, whereas the synthetic samples were kept in dry condition.

The specific loading protocol as depicted in Figure 1d was applied. After reaching a threshold force *f_t_* = 0.1 N, 10 preconditioning cycles were applied in the physiological loading regime with an amplitude *u_p_*= 8mm. This was followed by a displacement-controlled ramp loading until failure at a displacement rate of 1.3 mm/s. During the test, resulting displacement, force, applied strain, local samples strains, and where applicable, images, were acquired with a minimum sampling rate of 10 Hz. The full description of the loading protocol is provided in the supplementary material (section Appendix A, see Instructions_2nd_testround.pdf).

### 2.4. Data analysis

#### 2.4.1. Method variation

In addition to the testing data submission, in round 1 the participants were also requested to fill in a questionnaire to report their choices regarding pre-, intra-, and post-testing conditions. These included pre-testing storage conditions, test sample preparation and shape characterization, testing device characteristics, and testing protocol.

While the consensus protocol was used in round 2, the research groups were asked if they had any intentional or unintentional deviations from the protocol to identify potential sources of variability.

#### 2.4.2. Data submission

For each test, a table as shown in supplementary material was filled in, which differed slightly depending on the testing round. Apart from the aforementioned sample geometry information, actuator displacement, load cell values and local stretch estimation in the testing direction were obtained as a function of time. The participants were also asked to indicate whether the sample ruptured in the central area or at the clamp site.

#### 2.4.3. Data processing

##### Dimensions of interest

In round 2, the method and software used to measure dimensions from the calibrated images (see section 2.3.2) were left to the teams’ discretion.

##### Engineering stress & stretch

The raw force data was divided by the cross-section area of the reduced section to compute the raw engineering stress. The stretch data was retrieved by the participants using their own DIC analysis or marker tracking software. Subsequently, the raw stress and stretch data of all groups were processed by two data analysts to avoid any processing errors, before being synthesized by just one of them using Matlab 2019a and 2023a (Mathworks.com): only the test data between the end of preconditioning and the highest stress value was kept, filtered (forwardbackward moving average with a window of 5%) and resampled to 1000 points. A qualitative double-check of the cleaned data against the raw data was performed. Per step of 0.1 stretch ratio, a tangent modulus (TM) was computed using a least-squares line-fitting.

##### Comparison of stress/stretch curves

Various parameters for each team and per type of sample (i.e. a set) were computed to compare the results. The mean and standard deviation of the stress was computed at every 0.1 interval of the stretch, and the mean and standard deviation of the stretch was then computed at every 0.1 interval of the stress. Next, the mean stress/stretch curve a set of curves was computed point per point. I.e, for any given stress/stretch point of the 1000 points discretizing each curve, the mean of all the values at that point within the set was used.

##### Analytic uncertainty on stress

A theoretical uncertainty on the obtained stress values *U* (*σ*) was determined for each team and sample type, incorporating measurement uncertainties as follows:

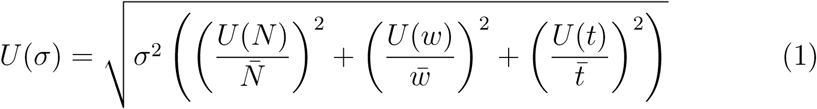

In the above equation, *σ* represents the engineering stress (in MPa), *N* is the normal force and *w* and *t* are the undeformed width and thickness of the sample, respectively. The measurement uncertainty on the force, *U* (*N*), is considered the same for all groups and on average assumed to be 0.1 N (based on a ISO 376 class 1 load cell measuring around 20 N), while the uncertainties on the width, *U* (*w*), and thickness, *U* (*t*), are both set to 0.02 mm (based on the maximum permissible error for external jaws on calipers with measurement length 0-5 cm, according to ISO 13385-1). The notation ·̅ represents the mean value of each parameter for a given team and sample type.

##### Zero-strain state

In the consensus protocol, the zero-strain state of the final upward ramp was considered to correspond to the time point *t_ref_* (see Figure 1d) at which the threshold force *t_f_* = 0.1 N was reached. However, during post-processing to increase consistency, this was modified to the strain state corresponding to the time point at which the measured stress was equal to 2∗*U* (*σ*) (see equation 1). This was equal to 0.03 MPa for biological samples and 0.06 MPa for synthetic ones.

## 3. Results

### 3.1. Submissions

In round 1, 20 participants successfully submitted their results for both the biological and synthetic data. In round 2, we received 17 successful submissions for the synthetic samples and 13 for the biological samples.

### 3.2. Method variation

The questionnaire results, regarding the choices of the research groups in round 1 for pre-, intra-, and post-testing conditions, are given in Figure 2 together with how many groups (in %) employed a certain technique or device.

**Figure 2:**
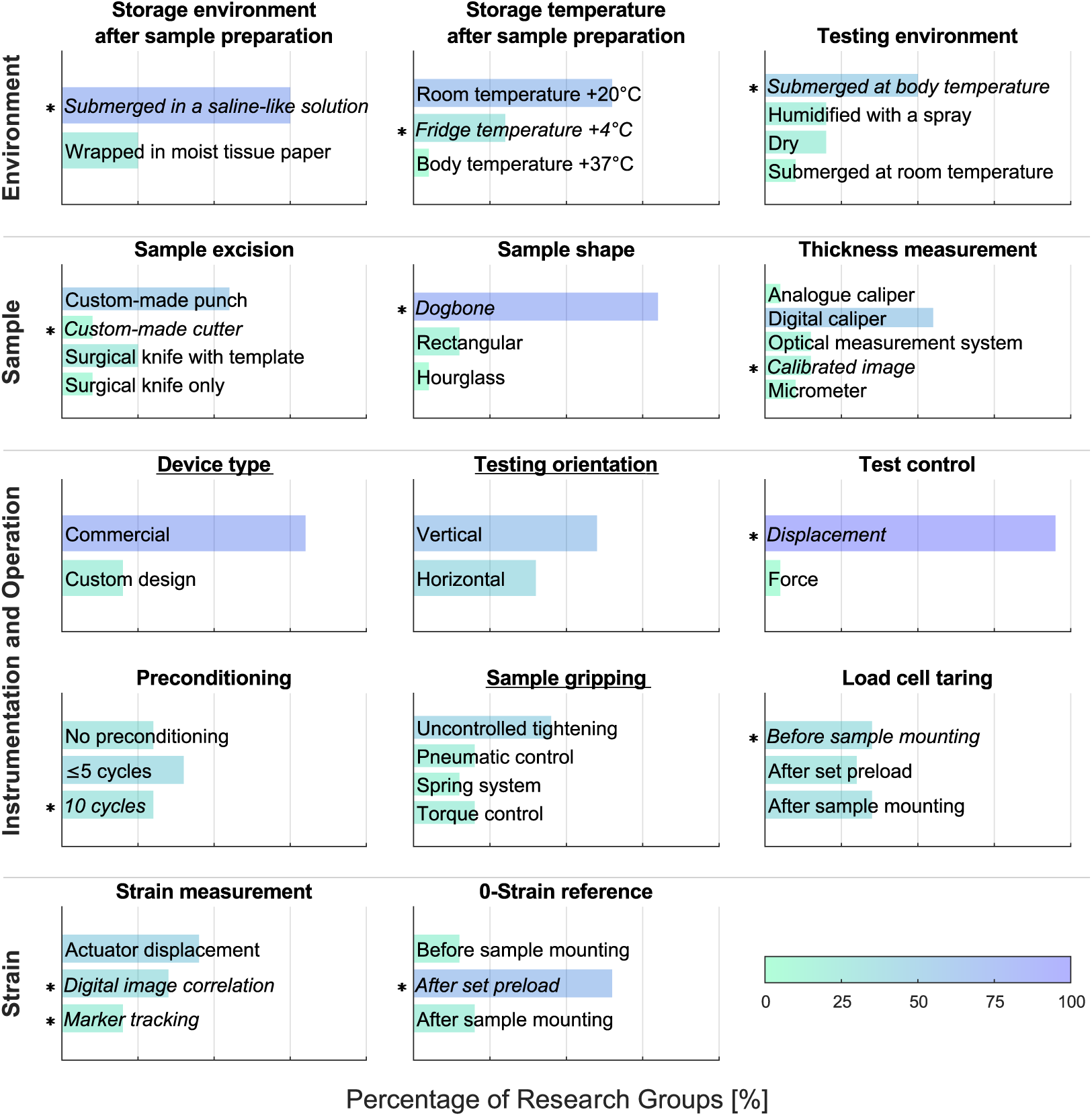
Method variation in round 1 and the percentage of groups using a certain approach. Parameters fixed in the consensus protocol of round 2 are marked with ‘∗’. Test criteria not constrained by the consensus protocol in round 2 are underlined.

Although the research groups were asked to use the consensus protocol in round 2, intentional or unintentional deviations from the protocol were observed by some groups and summarized in supplementary material (section Appendix A). For example, as the surface of the synthetic samples started showing cracks during testing, the stretch measurements could not be obtained with the intended DIC approach. Hence, many research groups reported the clamp-to-clamp distance instead. Another common deviation related to the removal of supposedly surrounding connective tissue in the biological samples. Some groups removed this tissue, possibly removing the adventitia along with it, others did not. Besides these, there were some occasional, unintentional deviations from the loading protocol, such as different preconditioning displacement or time point at which load-taring was performed.

### 3.3. Main results

An overview of all the engineering stress vs. stretch curves obtained by the participants for both testing rounds is given in Figure 3. These curves allow to obtain an impression of the overall variability in stress-stretch response and ultimate strain values of the tested samples.

**Figure 3:**
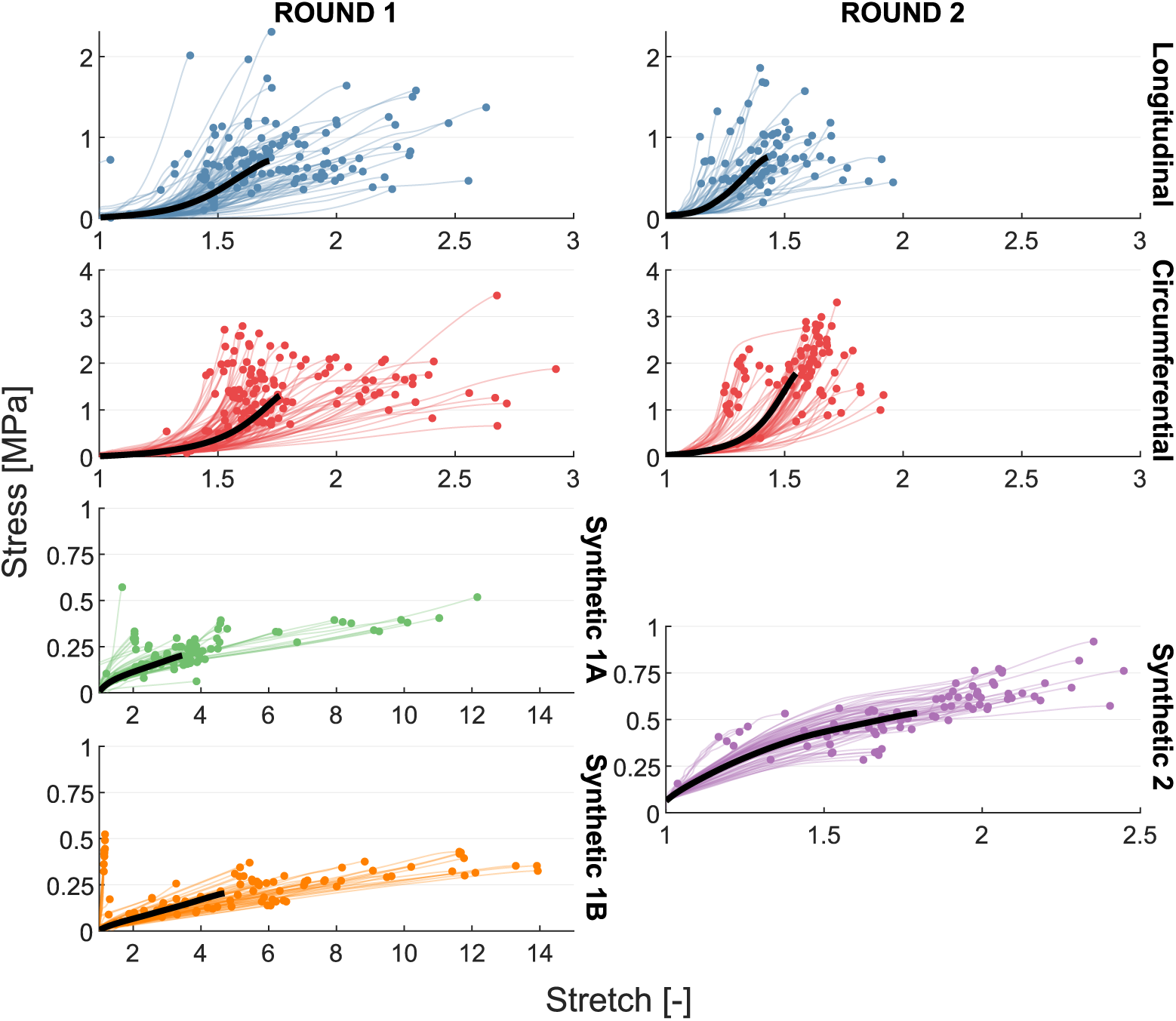
Engineering stress (in MPa) as a function of stretch obtained from uniaxial tension test until rupture, categorised by specimen type and separated per testing round. The average of all curves corresponding to each specimen type and testing round is represented in black.

Figures 4a and 4b show the stress vs. stretch curves obtained per research group on aortic tissue in the longitudinal direction during round 1 and 2 respectively. The corresponding curves for the circumferential direction and for the synthetic samples can be found in the supplementary material (Figure S4 and S5). The graphs also show the standard deviations of stress and stretch values at certain intervals. Figure 5 zooms in on this standard deviation and how it varies between research groups and between rounds. Now once again grouped for all participants, the graphs show the coefficient of variation of stress and stretch at two specific stretch levels. The graph also shows the percentage of research groups that fall under the arbitrary threshold of 0.33 in terms of this coefficient of variation.

**Figure 4:**
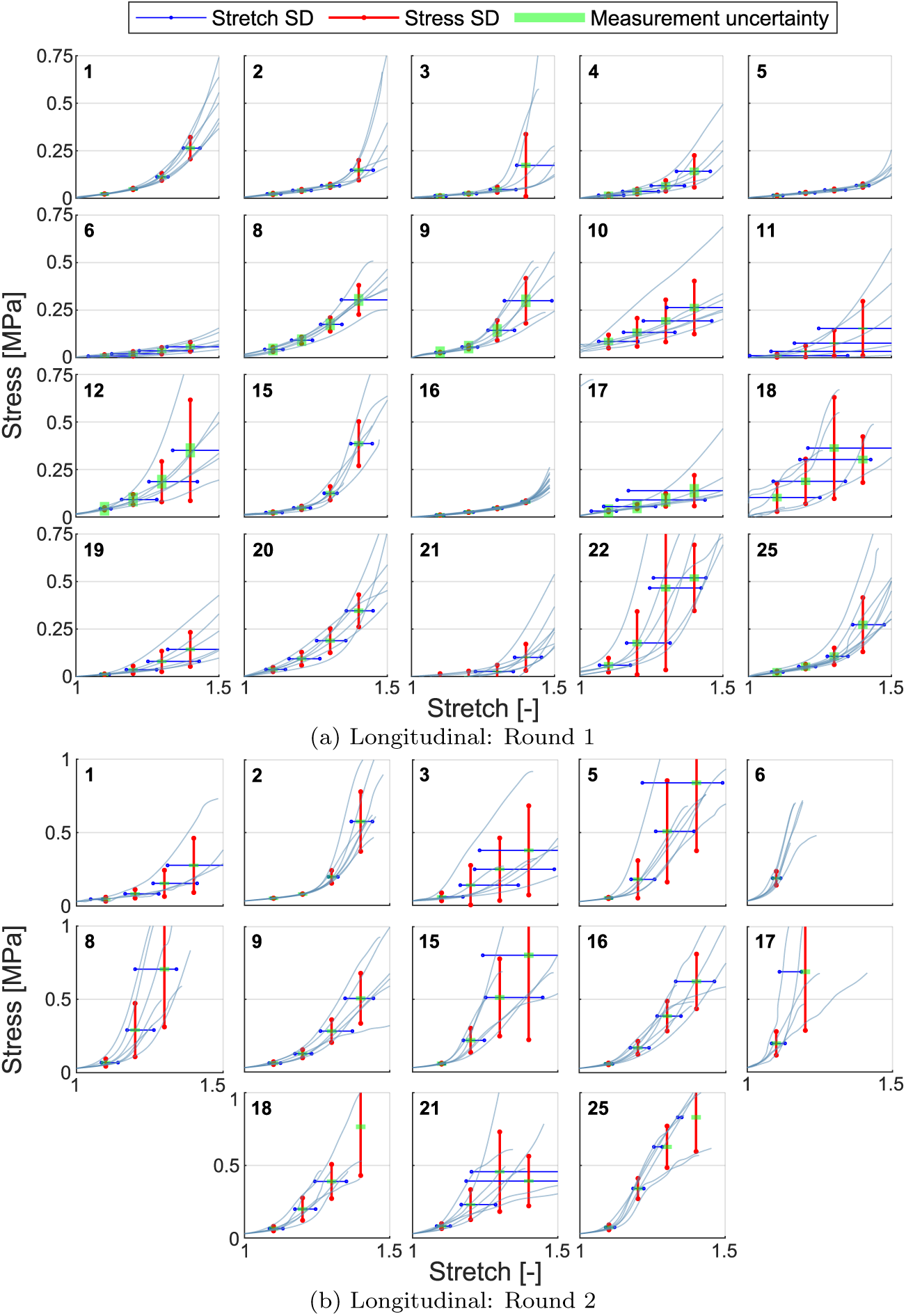
Engineering stress (in MPa) - stretch curves of longitudinal specimen types obtained in (a) round 1 and (b) round 2, grouped per participating research group indicated in the top left corner. Vertical lines represent the standard deviation (SD) of stress values at stretch levels of 1.1, 1.2, 1.3 and 1.4. Horizontal lines indicate the SD of stretch values at the mean stress value corresponding to these stretch levels. Measurement uncertainty on the mean stress value is shown at each stretch level, accounting for a 0.1 N load cell uncertainty and 0.02 mm uncertainty in specimen thickness and width measurements.

**Figure 5:**
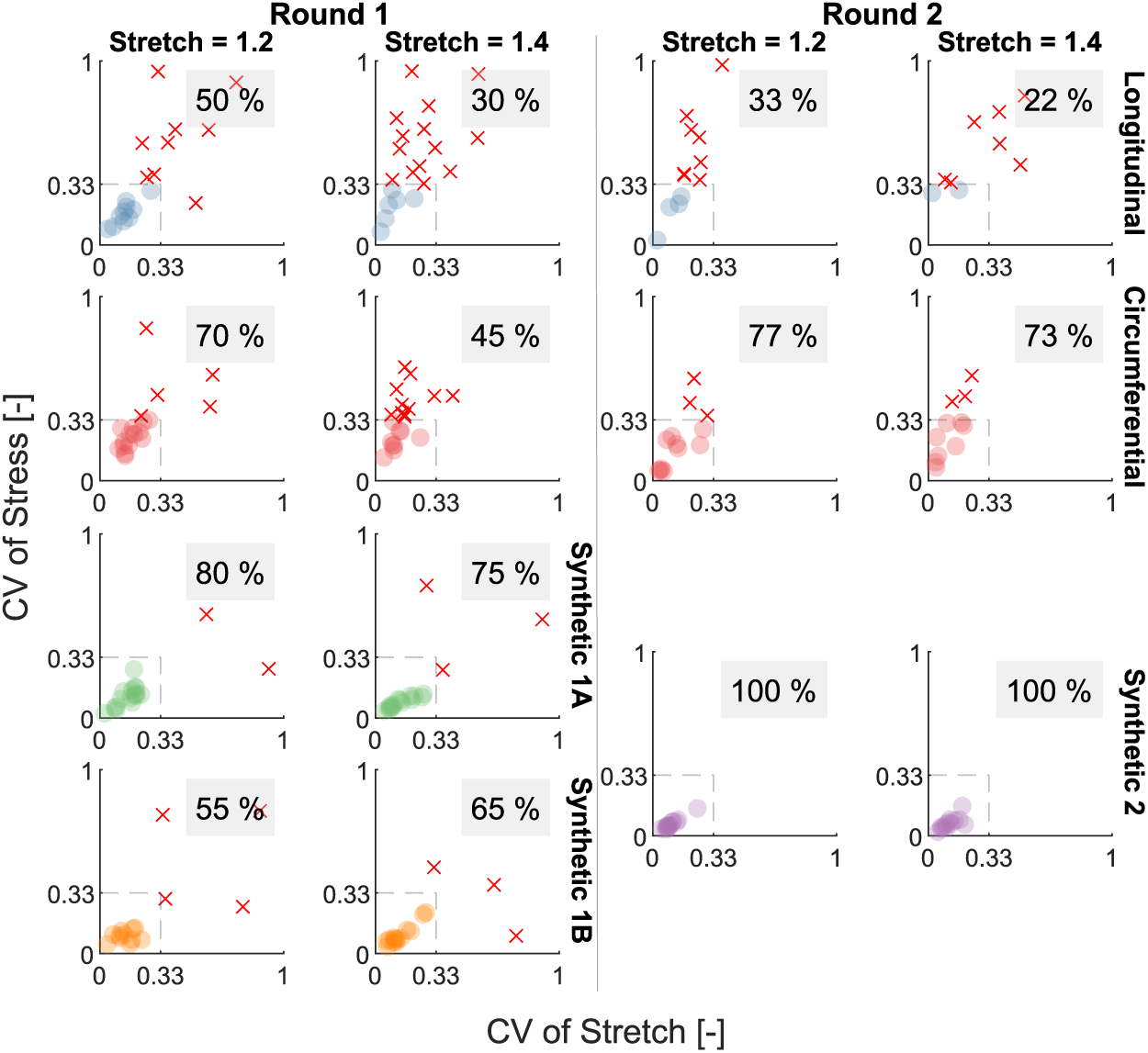
Coefficient of variation (CV) of engineering stress versus CV of stretch at two specific stretch levels, 1.2 and 1.4. CV values are calculated per research group. Research groups for which both CV values fall within the threshold of 0.33 are represented by dots, while those with either CV value exceeding this threshold are marked with ‘×’ markers. The percentage of research groups whose CV values remain within the threshold is indicated on each plot. The data are organised by specimen type and separated by testing round.

The distributions of the tangent moduli measured at 1.2 and 1.4 stretch are illustrated on Figure 6 and 7, respectively. To consider both the stress and strain measurement variabilities in the assessment of the tangent moduli, Figure 8a illustrates the coefficient of variation of stress and stretch at 1.2 and 1.4 stretch, for each group and for the various sample types. This figure discriminates the groups that fall under the arbitrary threshold of coefficient of variation of 0.33 for both stretch and stress.

**Figure 6:**
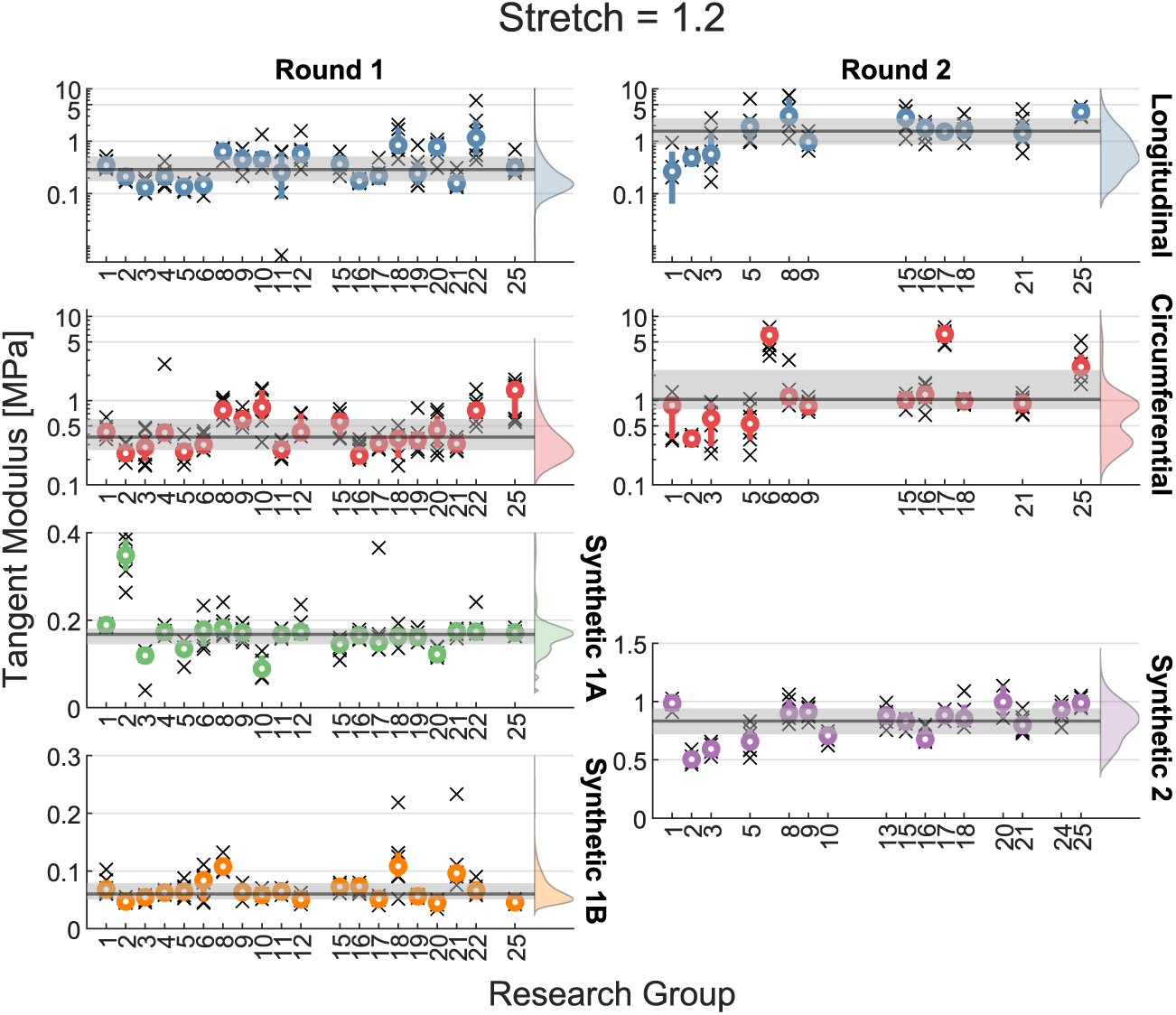
Tangent modulus of the stress-stretch curve at 1.2 stretch per research group, organised by testing round and specimen type. Individual moduli are represented by ‘×’ markers, while the mean for each research group is indicated by a dot and first- to third quantile (Q1-Q3) values are shown as bars. The overall mean and Q1–Q3 values, aggregated across all research groups, are depicted by a line and shaded band, respectively. A density plot is included to represent the distribution of all moduli for each specimen type and testing round, providing insights into the data distribution and variability.

**Figure 7:**
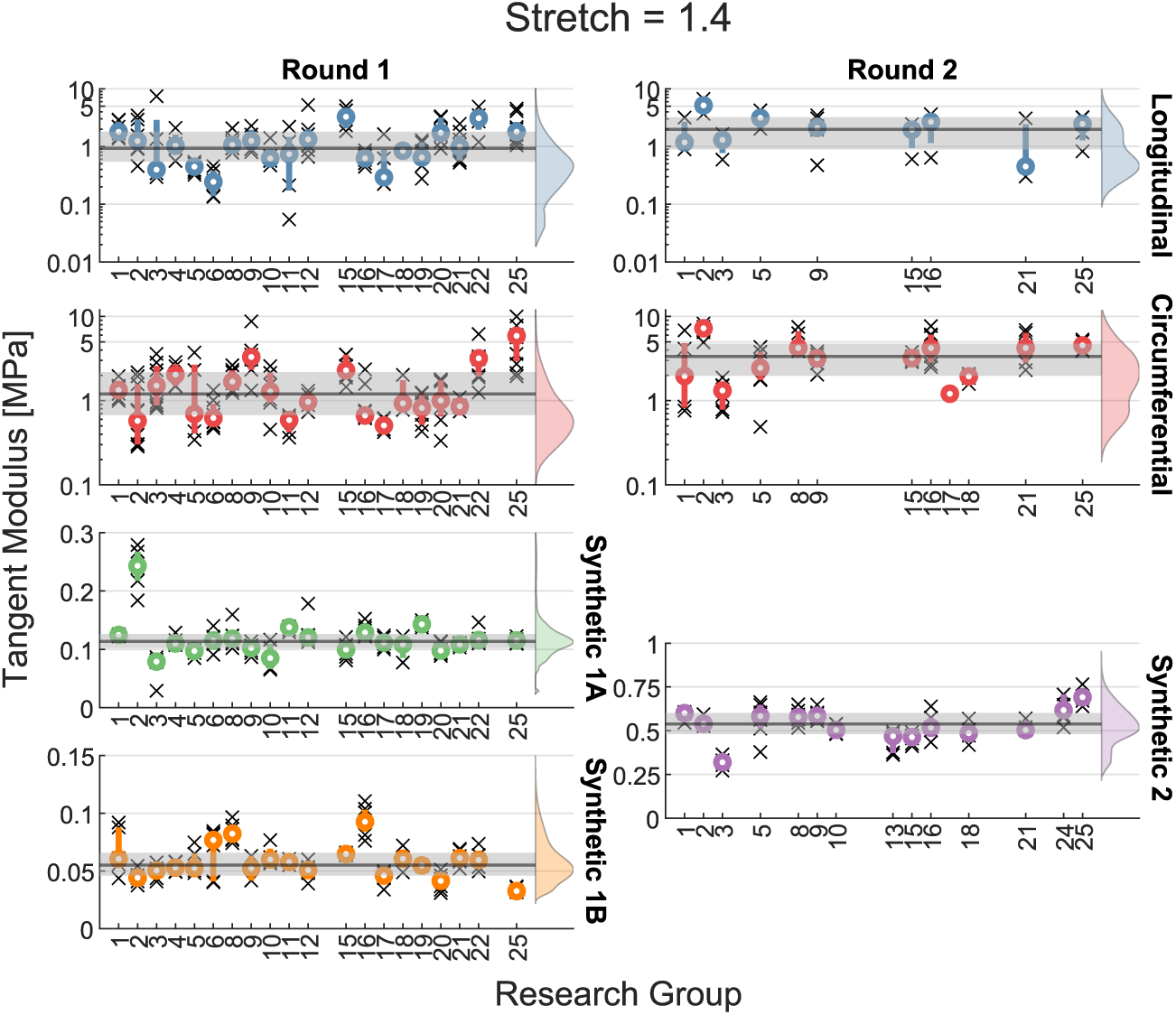
Tangent modulus of the stress-stretch curve at 1.4 stretch per research group, organised by testing round and specimen type. Individual moduli are represented by ‘×’ markers, while the mean for each research group is indicated by a dot and first- to third quantile (Q1-Q3) values are shown as bars. The overall mean and Q1–Q3 values, aggregated across all research groups, are depicted by a line and shaded band, respectively. A density plot is included to represent the distribution of all moduli for each specimen type and testing round, providing insights into the data distribution and variability.

**Figure 8:**
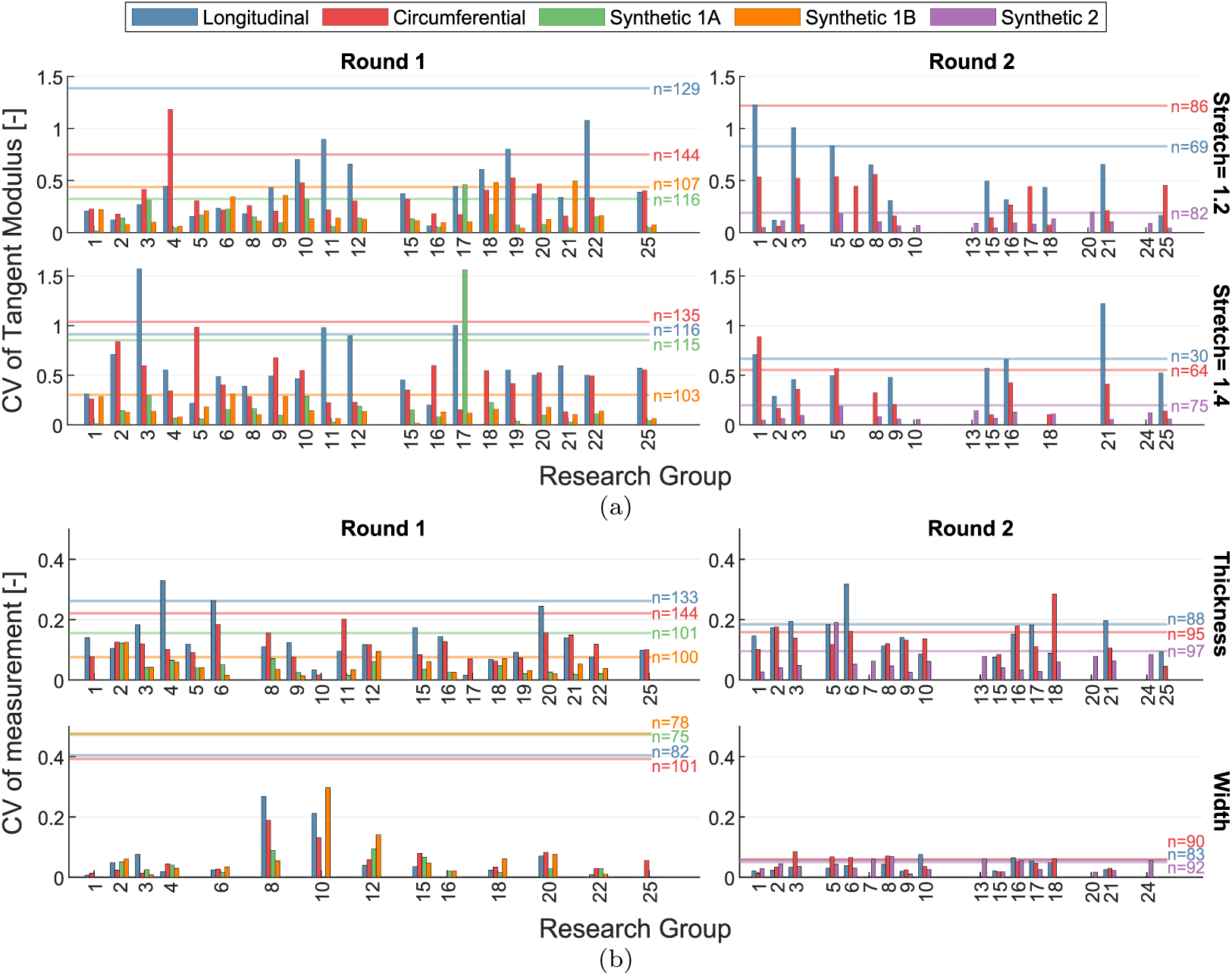
Bar plots showing the coefficient of variation (CV) of (a) the tangent modulus of the stress-stretch curve at 1.2 and 1.4 stretch values, and (b) thickness and width measurements. CV values are represented for each research group across different specimen types, separated by testing round. The overall CV value, calculated by aggregating measurements across all research groups for each specimen type, is represented by a line. The corresponding sample size is annotated adjacent to the line.

The distributions of the measurements of the thickness and the width of the samples among the groups and for both rounds are illustrated on Figure 9 and Figure 10, respectively. The coefficient of variation values of these measurements are illustrated on Figure 8b.

**Figure 9:**
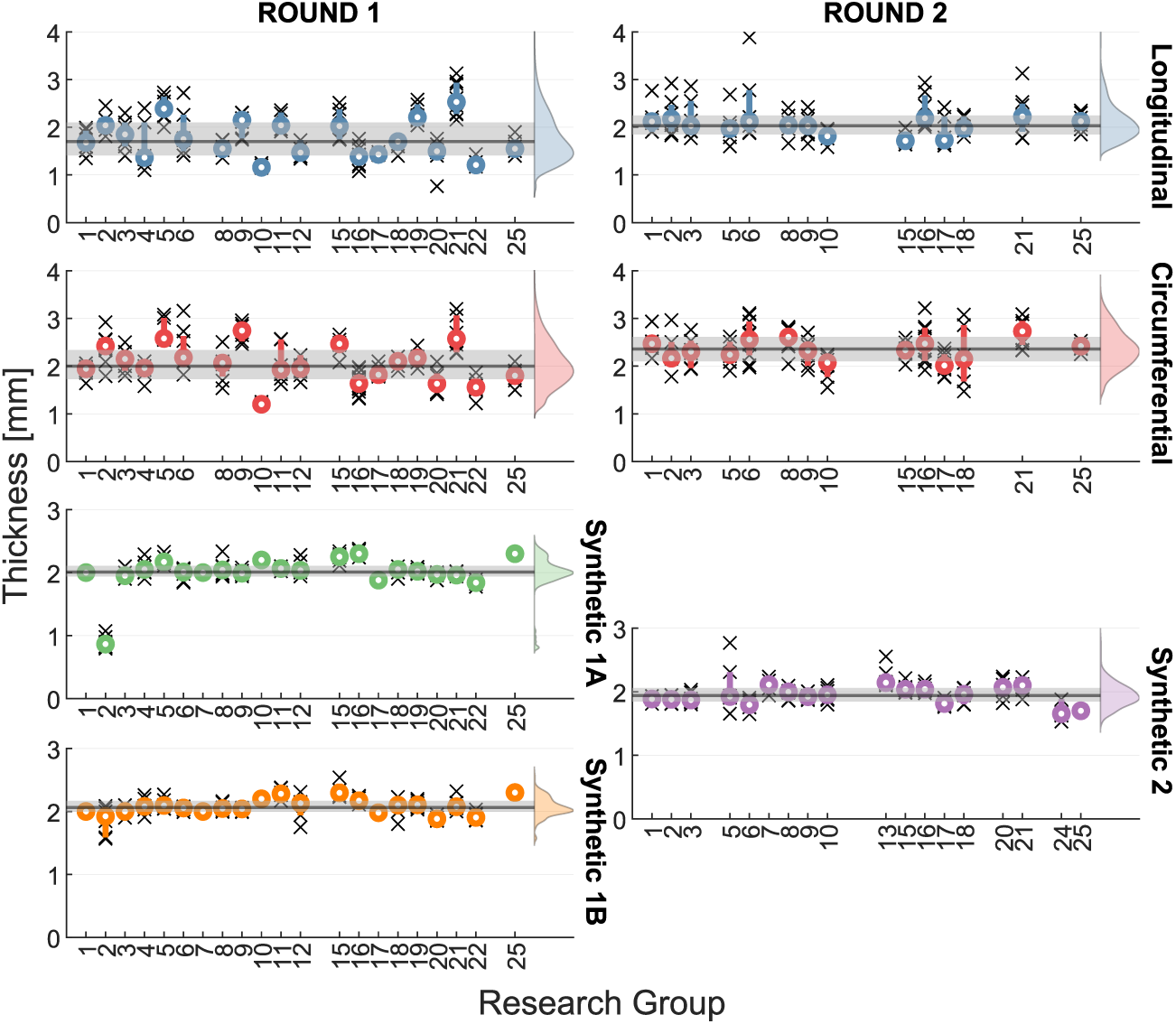
Specimen thickness measurements per research group, organised by testing round and specimen type. Individual measurements are represented by ‘×’ markers, while the mean for each research group is indicated by a dot and first- to third quantile (Q1-Q3) values are shown as bars. The overall mean and Q1–Q3 values, aggregated across all research groups, are depicted by a line and shaded band, respectively. A density plot is included to represent the distribution of all measurements for each specimen type and testing round, providing insights into the data distribution and variability.

**Figure 10:**
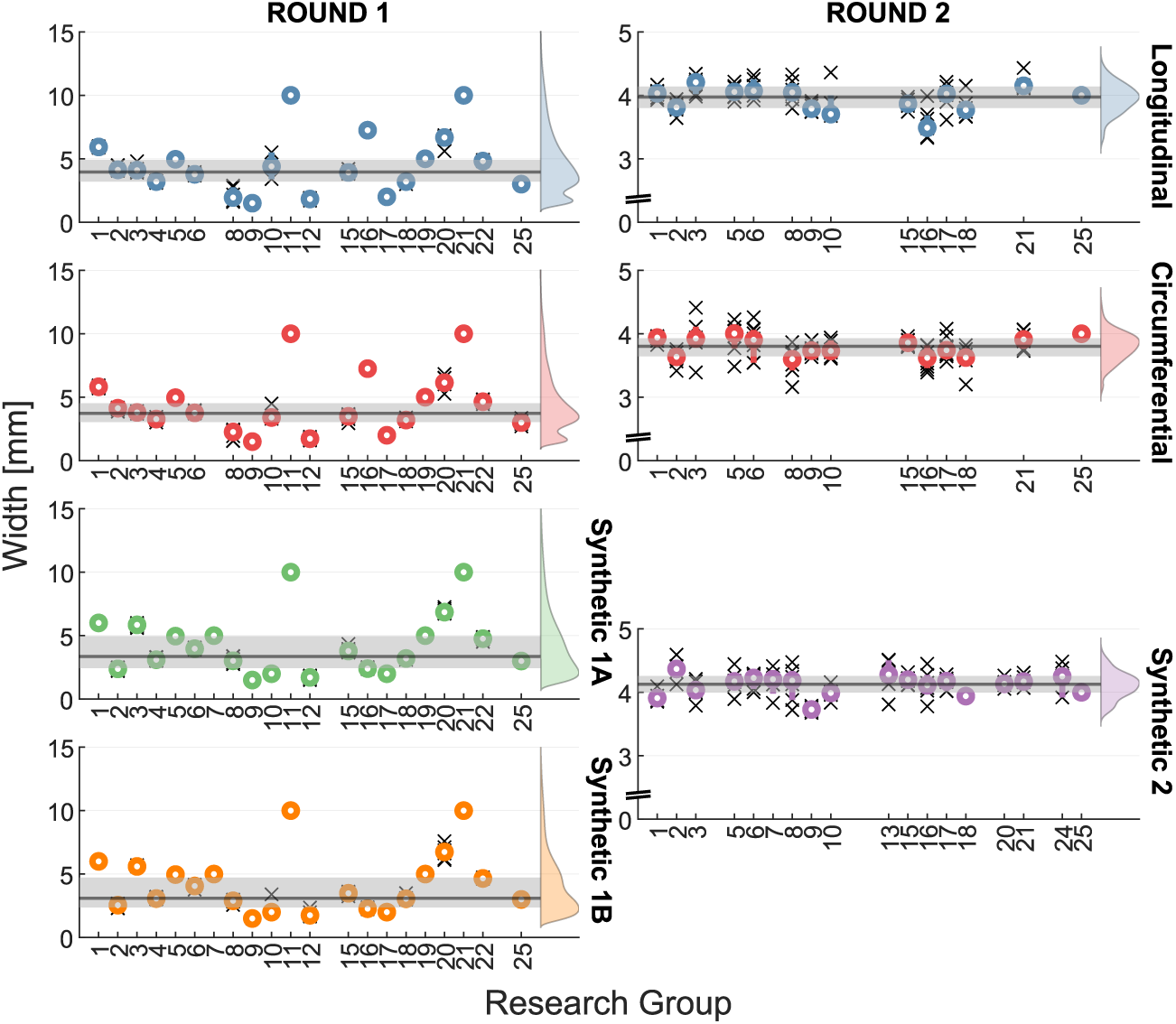
Specimen width measurements per research group, organised by testing round and specimen type. Individual measurements are represented by ‘×’ markers, while the mean for each research group is indicated by a dot and first- to third quantile (Q1-Q3) values are shown as bars. The overall mean and Q1–Q3 values, aggregated across all research groups, are depicted by a line and shaded band, respectively. A density plot is included to represent the distribution of all measurements for each specimen type and testing round, providing insights into the data distribution and variability.

## 4. Discussion

Even for a relatively simple method like uniaxial tensile testing, the results of round 1 demonstrate staggering variability between the participating laboratories. This is true for the obtained stress-stretch curves (Figure 3), the corresponding coefficient of variation values (Figure 5), as well as the resulting tangent moduli (Figure 8a). Looking at Figure 2, one can also observe a large variation in the choices made regarding the test protocol in this first round. However, statistical analysis (not shown here) could not identify any correlation between these protocol variations and the results. This implies that the observed variability cannot be directly attributed to a single queried aspect of the protocol. Indeed, the observed variability is a superposition of biological variability, methodological (or aleatory) uncertainty *and* methodological (or epistemic) discrepancy.

### 4.1. Variability versus uncertainty in round 1

To minimize biological variability, aorta samples were harvested from animals of the same age, weight and strain, as detailed in Section 2.2, and randomly distributed over the participants. However, even in such a population, biological variability is expected. Therefore, in an attempt to isolate the effect of biological variability, we also integrated synthetic samples into our analysis, as they suffer far less from intrinsic variability. Figure 8a shows how the overall coefficient of variation is indeed significantly reduced for the synthetic materials compared to the biological tissue, but still reaches levels close to 50%.

A rough estimate of the degree of methodological uncertainty on the stress measurement was made using equation 1, based on estimates of the accuracy of the used load cells and geometry measurement methods. Figure 4a shows how this estimated measurement uncertainty relates to the resulting standard deviation in stress. For most groups, this uncertainty represents only a fraction of the total observed variability, even for the synthetic material (see the supplementary material).

A significant portion of the observed variability must therefore be attributed to methodological variability, where we can in turn distinguish between intra-research group variability (also known as repeatability) and inter-research group variability (also known as reproducibility). Although one might expect the former to be minimal, given the instructions to test consistently, i.e. on the same day and by the same operator, the results indicate a significant and varying amount of intra-research group variability. Consequently, one might expect this repeatability to correlate with the reported protocol variations (see Figure 2). However, as mentioned, further statistical analysis did not reveal any correlation between these protocol variations and the actual results.

To disentangle sources of variability, one could attempt to obtain information on biological variability from other studies reported in literature. Indeed, there are several studies that report mechanical properties of aorta. For example, Shahbad et al. (2025) report on regional variations in stiffness of the human aorta along its length, whereas Ryu et al. (2022) report region-dependent stiffness parameters of porcine aorta. Both studies show a response and degree of variability (per region) in the same order of magnitude as our results. Of course, and this is the main message of our publication, it is not possible to distinguish methodological variability from biological variability in any of the studies. Indeed, there is no reason to assume that other laboratories would not suffer from the same degree of methodological variability as our reported intra-research group variabilities.

### 4.2. Consensus protocol and its trade-offs

Therefore, in round 2, we aimed to minimize methodological variability of as many aspects of the protocol as possible, leading to a lengthy consensus protocol. All participants also used the same cutting tool and thickness measurement tool in this second round. Still, in this quest for a harmonized protocol, we needed to consider two trade-offs. The first was a trade-off between methods that are most likely to yield biologically relevant results and methods with a lower complexity. The latter are more accessible for a broad group of researchers, and might also induce a lower amount of variability. As an example, it is easier to test samples in dried conditions than in physiological, submersed conditions. The latter requires mounting of a fluid bath, which makes it harder to mount the sample, and, depending on the orientation of the set-up, puts restrictions on the type of load cell. Although the dry state of the sample might not alter the intrinsic variability between samples compared to a submerged state, the extra requirements and manipulations could induce further methodological variability. Still, we opted for testing in immersed conditions. A counterexample in this respect is the fact that all tissue was frozen until the test day, rather than testing the tissue in fresh conditions. Whereas fresh tissue testing is, of course, more biologically relevant (Chow & Zhang, 2011; Stemper et al., 2007), freezing the tissue was necessary to allow for uniformity in the tissue preservation step. The second trade-off was to be made between methods that are likely to yield the most unbiased results and methods that were available to most participants (in terms of infrastructure and personnel requirements). As an example, the thickness measurement method should ideally not compress the tissue (Kim et al., 2011; O’Leary et al., 2013) or correct for any compression during the measurement (Schwarz et al., 2023), but requires specific hardware and software that was not available to all participants, which is why we opted for the supplied thickness measurement tool.

ASTM D412 is a common standard to characterize mechanical properties of rubbers and elastomers using a tensile test. Our resulting consensus protocol deviates from this standard in various ways. Firstly, we used a lower displacement rate (1.3 mm/s vs. 500 mm/min) to approximate ‘physiological’ loading conditions. Secondly, where the standard suggests a fixed temperature, we allowed variations between laboratories in ambient temperature when testing the synthetic samples, which might contribute to the variability of these measurements. Due to the limited size of an aorta, our chosen dogbone shape was smaller than the standard, especially regarding the width of the grip section (8.1mm vs. 25mm) and the length of the reduced section (16mm vs. 33mm). This reduced size and aspect ratio might lead to a non uniform strain field within the area of measurement (Lin et al., 2024).

### 4.3. Variability on geometrical measurements

Despite efforts to reach a consensus on the testing methodology, the second round did not display significant improvement in the quantities of interest. One metric that was however drastically improved was the variation on the sample width measurement (see Figures 8b and 10). This makes sense given the fact that this dimension was not specified in the first round, whereas all participants used the same cutting tool in the second round. Thickness measurement variability was also improved (as can be observed from the reduced coefficient of variation of the thickness measurement in Figure 8b), but not to the same extent as the width measurement. This may be explained by a greater variation in the thickness compared to the width of the samples, as the samples were prepared using the cutting tool. Of course, release of residual stresses may still have led to some width variation in the samples. Secondly, the samples are approximately twice as wide as they are thick, reducing the signal to noise ratio for the latter measurement. The variability on the thickness measurement is indeed significantly lower for the synthetic samples in both rounds, although one can notice an outlier group for the measurements on synthetic 1A samples. Apparently, this group had used an optical laser measurement method, which suffered from laser light penetration for this sample type, yielding a systematic error in their results (O’Leary et al., 2013).

### 4.4. Variability on stress-stretch curves and tangent modulus

Figures 5 and 8a illustrate how the improvements on the dimension measurements did not propagate to the resulting stress-stretch curves or tangent moduli. For low stretch values, coefficient of variation levels are even slightly increased, and there is a decrease in research groups reaching the threshold value of coefficient of variation for the longitudinal stress-strain behaviour of the aorta. For the higher strain level, we do observe a reduction in coefficient of variation levels and a higher number of participants reaching the threshold, but the improvement is far lower than initially anticipated.

An explanation for this lack of improvement in terms of tangent modulus variability could lie in the following: due to the standardization of the zero strain state in the consensus protocol, the origin of the curves is more homogeneous and the curves have therefore on average shifted towards the left, leading to higher stresses for a certain stretch level (see Figures 6 and 7). As a consequence, the values of the tangent moduli obtained in round 2 are significantly higher than in round 1, increasing the amplitude of their variability.

In all cases, we can observe less variability for synthetic samples than for aortic samples. In round 2, all participants are below the coefficient of variation threshold value for the stress-stretch behaviour and the overall coefficient of variation in tangent modulus is lower for both stretch levels. The resulting tangent modulus is also significantly higher for the synthetic material used in round 2 compared to those of round 1. Indeed, a different material was used, which at least at low stretch levels, matched the average tangent modulus of the aortic material more closely.

### 4.5. Remaining sources of variability

The estimated methodological uncertainty on the stress measurement (equation 1) is significantly smaller than the observed one, even more so for round 2 (see Figure 4b). This means that much of the variability is still due to other factors, a number of which are discussed below.

- **Stretch measurement uncertainty** - Either marker tracking or DIC was performed in round 2. The reliability of the former can be affected by marker placement accuracy and tracking resolution, which was not harmonized between the participants. Also for DIC, its reliability depends on the quality of the speckle pattern, the resolution of the imaging system, as well as on software analysis settings, as was investigated at length in a series of ‘DIC challenges’ (Reu et al., 2018, 2022). Nevertheless, these DIC challenges were not specifically aimed at biological tissue applications, which pose particular challenges. For example, immersion of the sample in a water bath possibly causes optical distortion, speckle pattern application should ensure properly adherence to the tissue, not cause or require dehydration of the tissue, and last through large deformations (Lionello et al., 2014). The sample shape, shorter than the ASTM recommendations for synthetic materials, can induce an inhomogeneous deformation pattern, the degree of which should be investigated (Lin et al., 2024). These inhomogeneities should be dealt with, either through a sample-size specific correction factor, or by applying DIC rather than marker tracking where possible.
- **Zero-strain state** - The samples were considered to be in their zero strain state at the start of the test, specifically at 0.03 MPa for biological samples and 0.06 MPa for synthetic ones. This stress-based definition clearly induces error as well as uncertainty proportional to the force and surface area measurement uncertainty. Moreover, it is generally know that biological tissues exhibit residual strains (Vaishnav & Vossoughi, 1983), further confounding the definition of the zero-strain state.
- **Adherence to the testing protocol in round 2** - Despite the provided instructions, we observed variations in how different groups adhered to the testing protocol. For instance, some groups removed the connective tissue of the adventitial layer while others did not. The adventitial layer is known to be collagen-rich and therefore contribute significantly to the nonlinearity of the stress-stretch curve. Additionally, there were discrepancies in the applied preconditioning cycles and in the definition of the state to be reached before starting the final ramp loading. This might be attributed to misunderstanding of the instructions, but also due to the incompatibility of certain testing machine controllers w.r.t. the desired protocol. These deviations are documented in detail in the supplementary material (see ProtocolDeviationsRound2.pdf). Note also that the hypothesis was made that each participant used calibrated sensors and testing machines, for example according to ASTM E4 standards, but this was not explicitly verified.
- **Intra-group variability** - Some teams exhibited greater intra-group variability in round 2 compared to round 1. This could be attributed to the fact that they were following a testing protocol different from their usual practices. For example, managing samples in an immersed test environment can be more challenging and may introduce additional variability, especially when the user is still going through a learning curve.
- **Sample misalignment** - For highly anisotropic samples, ensuring that samples are cut precisely along the circumferential or longitudinal direction is crucial for consistent results. Misalignment can lead to variations in the mechanical properties measured, as the orientation of the fibers in the tissue can significantly affect its behavior. Careful attention to cutting and aligning the samples is necessary to minimize this source of variability, but can never be fully excluded.
- **Variability of the synthetic materials** - The synthetic samples were considered as isotropic and homogeneous, although it is known that the manufacturing process may introduce a flow-related anisotropy and other imperfections. Hence, even for these materials, it is important to differentiate methodological variability from production-related variability.
- **Participant preferences** - Certain aspects of the testing protocol were left to the discretion of the participants, such as the orientation of the testing device (vertical vs horizontal), the type of clamps used (pneumatic or manual). Narrowing this down would have led to a much smaller number of participants, which is why this trade-off was made. Scholze et al. (2020) have proposed an easy-to-fabricate clamping system for uniaxial tensile testing, that could be investigated for use in future testing rounds.
- **Number of test samples** - 6 samples were tested per condition per research group, each time coming from two different animals. This number is based on previous experiments combined with the practical limitations on the number of aortas that could be collected from the slaughterhouse. Ideally, a dedicated power analysis of the current results should be performed to reveal the required number of tests for future testing rounds for this specific test and sample type.

### 4.6. Future work

This pilot campaign serves as an exploration of the current landscape, highlighting the existing variability in the mechanical testing to characterize biological tissues. As such, it marks only the start of a much needed effort to further quantify and minimize this variability, and the following areas of future work have been identified.

- Further statistical analysis of the data should be performed to help identify specific sources of variability. Also, even more detailed analysis of each of the participants’ submissions might uncover further deviations from the protocol.
- Each of the above listed remaining sources of variability accounts for just a portion of the overall variability. For example, when observing the degree of anisotropy of the biological tissues tested here, one could argue that the difference between the circumferential and longitudinal behavior is much smaller than the overall variability, such that sample misalignment isn’t such a big issue. Indeed, the challenge is not only to identify sources of uncertainty, but also to rank each of these sources in terms of their relative importance. Once a clear overview and ranking of these sources of variability is obtained, further fine-tuning of the consensus protocol can be performed after which a new test round can be launched.
- To assess the impact of operator variability, it would be beneficial to have the same operator perform the test on different set-ups. In other words, rather than sending samples around, one could exchange researchers. Organising training sessions or workshops for all participants could help improve consistency in following the testing protocols.
- The shape and amplitude of the stress-stretch curves of all the synthetic materials deviate strongly from those of the curves for biological samples. This severely limits the relevance of these synthetic samples in terms of protocol verification capacity. Indeed, these synthetic samples do not exhibit the characteristic nonlinear stiffening behaviour, and this absence of a so-called ‘toe-region’ significantly reduces the sensitivity to slight differences in strain. Therefore, for future testing rounds, we need synthetic materials that do exhibit this nonlinear stiffening behaviour. Possible candidates are PVA hydrogels (Millon et al., 2006), fabric-reinforced polymers (Zhalmuratova et al., 2019) or melt electrowritten scaffolds (Mirani et al., 2024). A thorough examination of these materials is required to ensure maximal resemblance to aortic tissue with minimal variability due to the production process, storage, etc.
- To generalize our findings it is necessary to expand the study to include other types of tests and tissues. This can help to identify whether the observed variability is specific to certain tests or tissues, or if it is a more general issue.

This first C^4^Bio campaign serves as a wake-up call that current methodological variability is too large to enable reliable comparison of results between research groups. This undermines the credibility of resulting material properties reported in literature, and their consequent use in *in silico* simulations. This study has made it clear that proper uncertainty quantification is essential, and that through community effort we should aim to further increase the quality of our mechanical testing methods and reduce the uncertainty to a workable level.

## 5. Acknowledgements

The authors would like to thank the Avicenna Alliance and VPH Institute for their financial support, as well as KU Leuven (OPSTAMET C2M/24/031 and KERNFAC FIBER) and the Research Foundation - Flanders (I011822N and G0H1116N). Furthermore, we wish to acknowledge the efforts of the staff of FIBEr, KU Leuven core facility, specifically Adriana Wilmots Helsen, Markos Kapeliotis and Wouter Willekens, for procuring, preparing and shipping all samples. Next, we would like to express our gratitude to Eddy Smets and Thomas Pilkington from the Fablab of Mechanical Engineering, KU Leuven for their assistance in the development of the cutting tool used in test round 2. Finally, we would also like to thank the following researchers for their contribution to the conception of C^4^Bio, to the mechanical testing and/or to the data-analysis:

- Dr.Ir. Klaas Vander Linden, KU Leuven
- Dr. Kris Bogaerts, KU Leuven
- Ir. Thibault Vervenne, KU Leuven
- Ir. Matijs Loeckx, KU Leuven
- Dr. Alex Caulk, Medtronic
- Dr. Markus Reiterer, Medtronic
- Dr. Venkateswaran Perumal, Stryker
- Dr. Raja Jayendiran, Stryker
- Dr. Naveen Kumar Subramani, Stryker
- Dr. Martin Tanaka, University of Western Carolina
- Dr. Connor Gains, Queen Mary University of London

## Appendix A. Supplementary material

All the supplementary material can be found in a repository on Zenodo through the following link.

https://zenodo.org/uploads/14179432?token=eyJhbGciOiJIUzUxMiJ9.eyJpZCI6IjRiMzM5MGIyLTNlYTItNGY4NS04ZTliLWFhMjA2ZTMzN2QyNiIsImRhdGEiOnt9LCJyYW5kb20iOiJjMGZmZDUyZWNlZDIzYzc3MTYwNjAyMmRlNjZhYzAwNyJ9.PG4xkyAfg2JTQuDuZfFh341z2F2nWUedUd9D33ey0aP1AD4s4XTtyyJpmtbftX14oKjP4kSFhu8GZC4ZiJhRfA

*Note to reviewers: This link will be shortened in the final submission, once the repository is published*.

