## Supplementary materials for "Community Challenge towards Consensus on Characterization of Biological Tissue: C^4^Bio’s First Findings"

### 1. Materials & methods

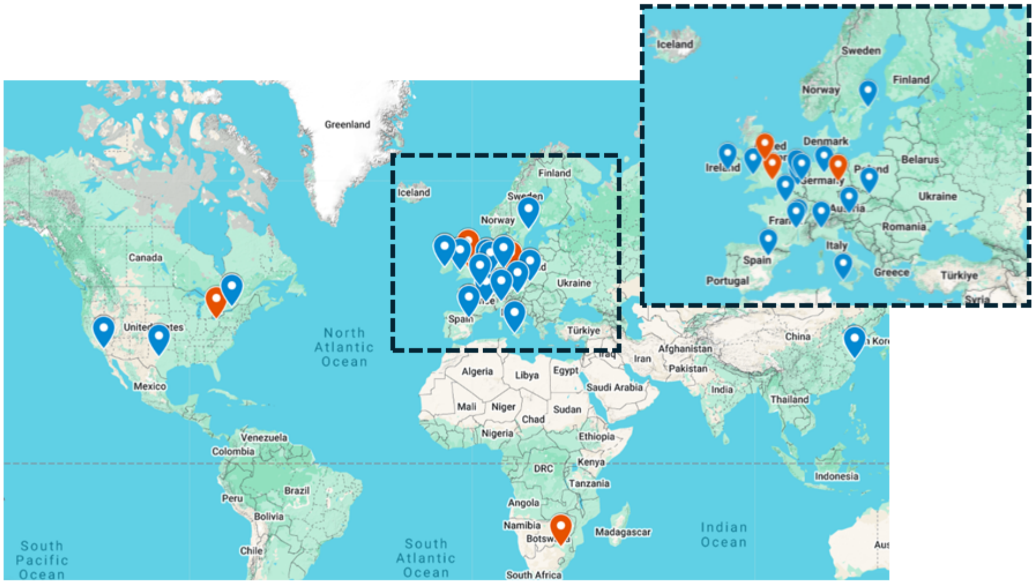


Figure S1: Overview of the participating research groups of the first (red+blue) and second (blue) testing round.

#### 1.1. *Sample preparation & shipping*

##### 1.1.1. *Biological tissue*

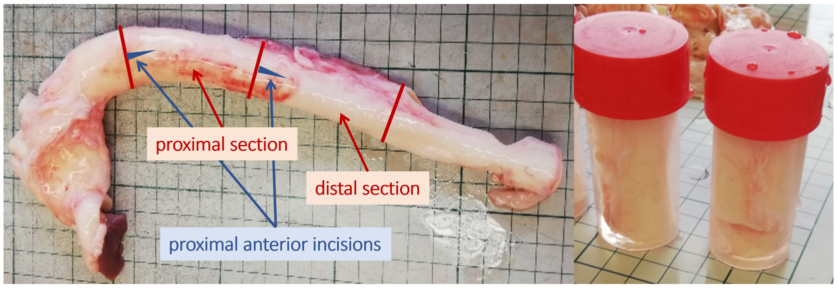


Figure S2: Full aortic arches were divided into a proximal and a distal section of approximately 5 cm in length and placed in separate containers in physiological serum.

##### 1.1.2. *Synthetic samples*

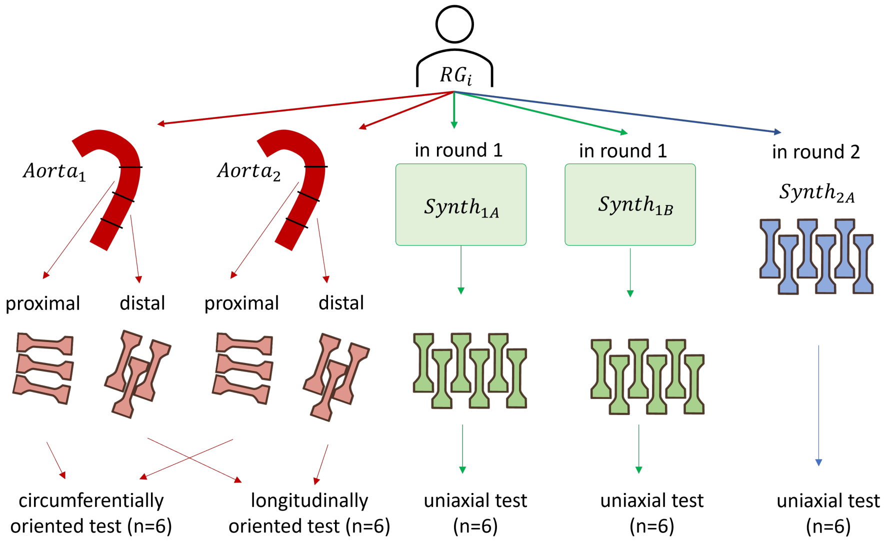


Figure S3: Overview of the samples tested per participating research group RGi in both testing rounds. Participants cut their own samples from the received segments, except for the synthetic material of round 2, which was 3D printed into the desired shape.

#### Standard Deviation on Stress/Stretch

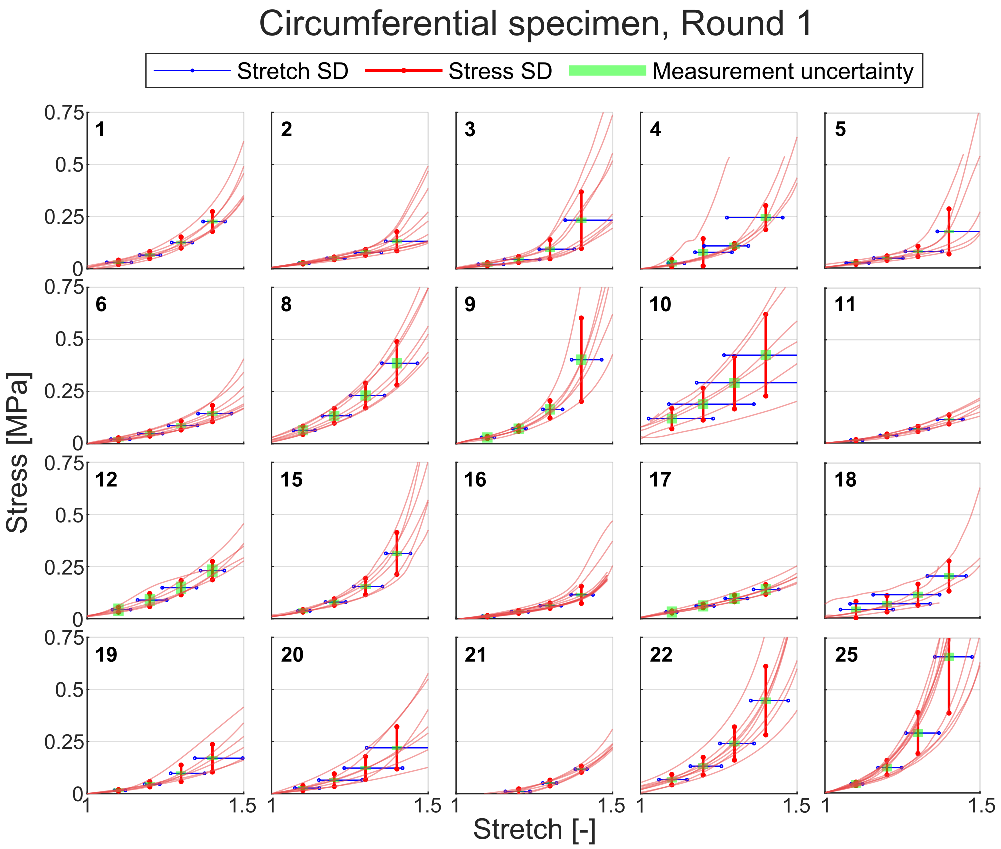


Figure S4: Engineering stress (in MPa) - stretch curves of circumferential specimen types obtained in round 1, grouped per participating research group indicated in the top left corner. Vertical lines represent the standard deviation (SD) of stress values at stretch levels of 1.1, 1.2, 1.3 and 1.4. Horizontal lines indicate the SD of stretch values at the mean stress value corresponding to these stretch levels. Measurement uncertainty on the mean stress value is shown at each stretch level, accounting for a 0.1 N load cell uncertainty and 0.02 mm uncertainty in specimen thickness and width measurements.

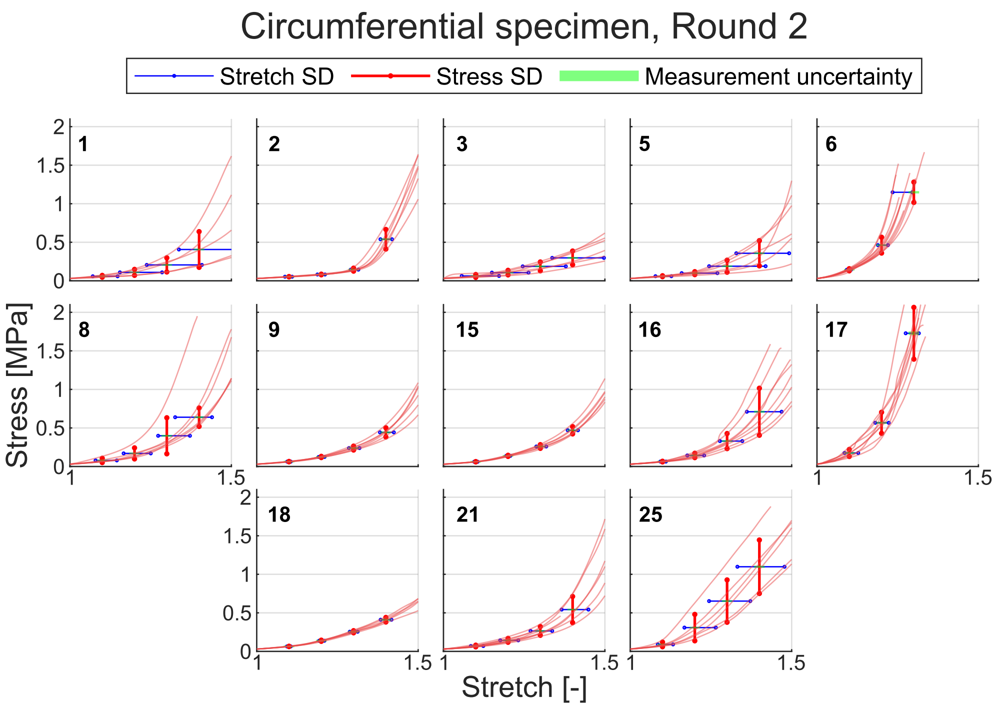


Figure S5: Engineering stress (in MPa) - stretch curves of circumferential specimen types obtained in round 2, grouped per participating research group indicated in the top left corner. Vertical lines represent the standard deviation (SD) of stress values at stretch levels of 1.1, 1.2, 1.3 and 1.4. Horizontal lines indicate the SD of stretch values at the mean stress value corresponding to these stretch levels. Measurement uncertainty on the mean stress value is shown at each stretch level, accounting for a 0.1 N load cell uncertainty and 0.02 mm uncertainty in specimen thickness and width measurements.

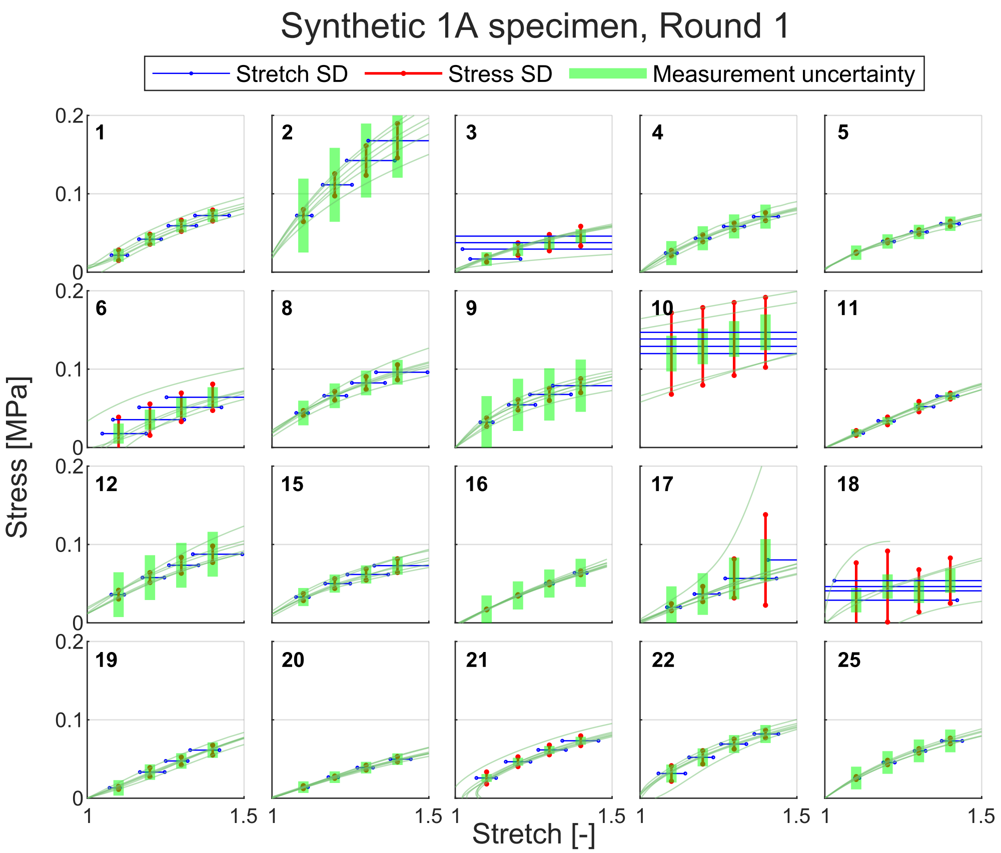


Figure S6: Engineering stress (in MPa) - stretch curves of synthetic 1A specimen types obtained in round 1, grouped per participating research group indicated in the top left corner. Vertical lines represent the standard deviation (SD) of stress values at stretch levels of 1.1, 1.2, 1.3 and 1.4. Horizontal lines indicate the SD of stretch values at the mean stress value corresponding to these stretch levels. Measurement uncertainty on the mean stress value is shown at each stretch level, accounting for a 0.1 N load cell uncertainty and 0.02 mm uncertainty in specimen thickness and width measurements.

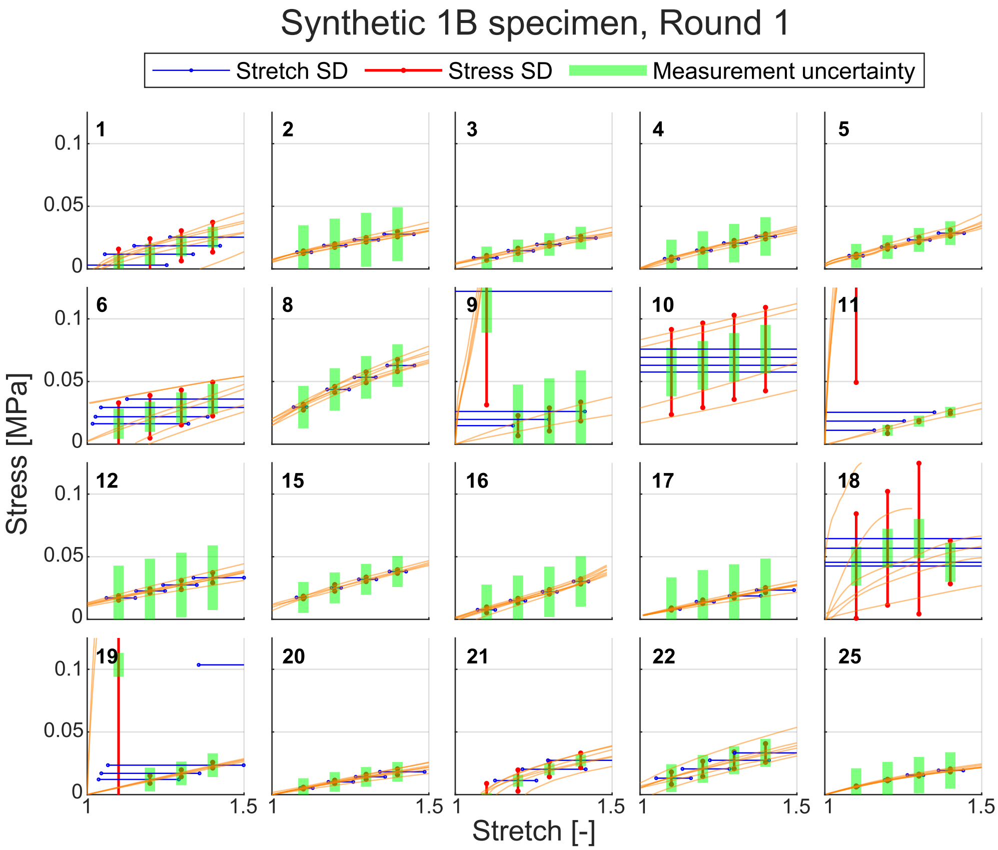


Figure S7: Engineering stress (in MPa) - stretch curves of synthetic 1B specimen types obtained in round 1, grouped per participating research group indicated in the top left corner. Vertical lines represent the standard deviation (SD) of stress values at stretch levels of 1.1, 1.2, 1.3 and 1.4. Horizontal lines indicate the SD of stretch values at the mean stress value corresponding to these stretch levels. Measurement uncertainty on the mean stress value is shown at each stretch level, accounting for a 0.1 N load cell uncertainty and 0.02 mm uncertainty in specimen thickness and width measurements.

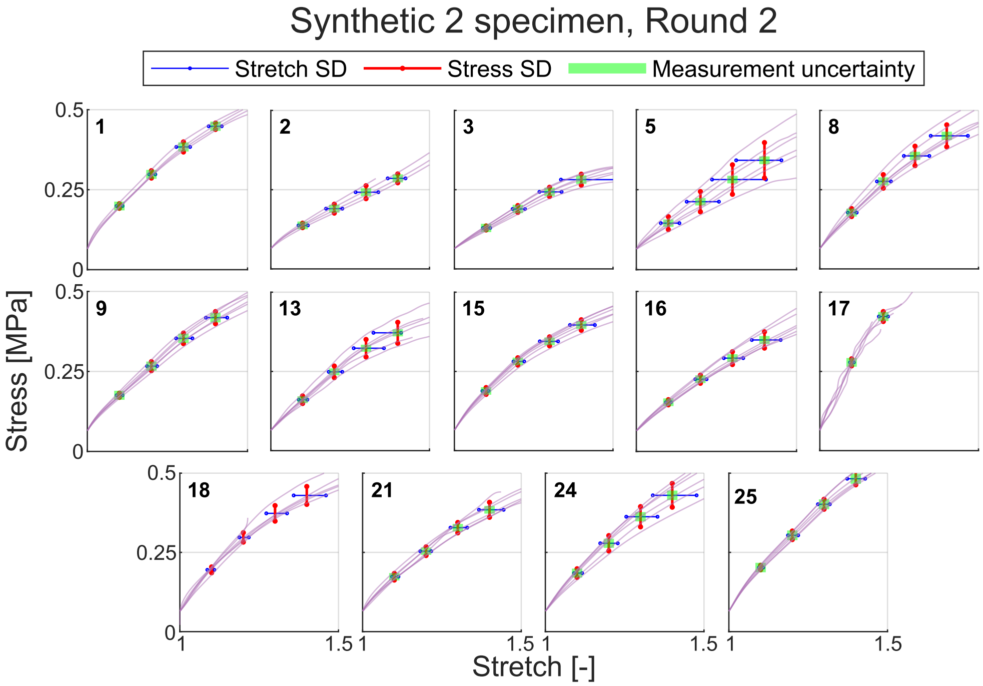


Figure S8: Engineering stress (in MPa) - stretch curves of synthetic 2 specimen types obtained in round 2, grouped per participating research group indicated in the top left corner. Vertical lines represent the standard deviation (SD) of stress values at stretch levels of 1.1, 1.2, 1.3 and 1.4. Horizontal lines indicate the SD of stretch values at the mean stress value corresponding to these stretch levels. Measurement uncertainty on the mean stress value is shown at each stretch level, accounting for a 0.1 N load cell uncertainty and 0.02 mm uncertainty in specimen thickness and width measurements.

#### Coefficient of Variation of Stress/Stretch

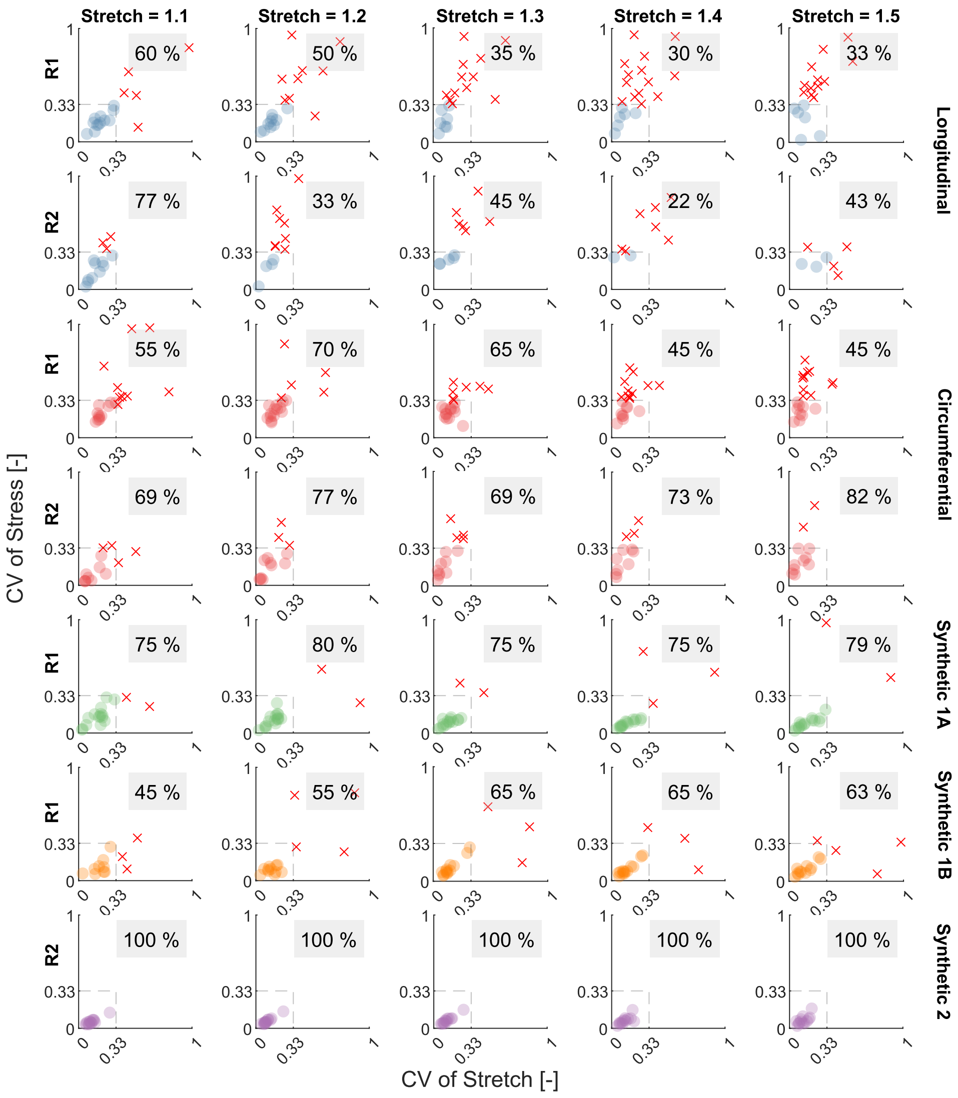


Figure S9: Coefficient of Variation (CV) of engineering stress versus CV of engineering strain at stretch levels 1.1 to 1.5 with 0.1 steps. CV values are calculated per research group. Research groups for which both CV values fall within the threshold of 0.33 are represented by dots, while those with either CV value exceeding this threshold are marked with '×' markers. The percentage of research groups whose CV values remain within the threshold is indicated on each plot. The data are organised by specimen type and separated by testing round 1 (R1) and 2 (R2).

#### Tangent Modulus (TM)

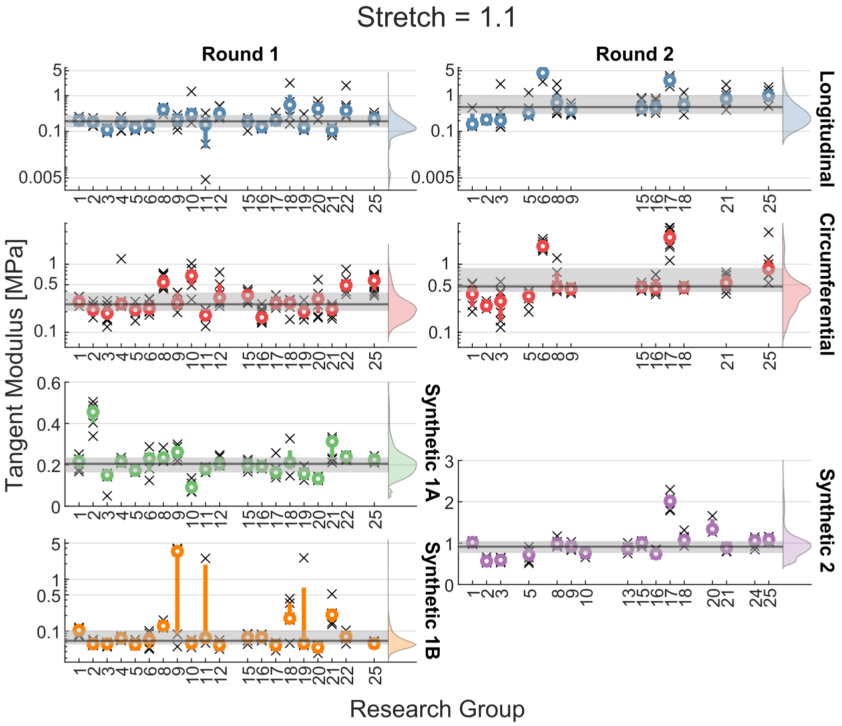


Figure S10: Tangent modulus of the stress-stretch curve at 1.1 stretch per research group, organized by testing round and specimen type. Individual moduli are represented by '×' markers, while the mean for each research group is indicated by a dot and first- to third quantile (Q1-Q3) values are shown as bars. The overall mean and Q1–Q3 values, aggregated across all research groups, are depicted by a line and shaded band, respectively. A density plot is included to represent the distribution of all moduli for each specimen type and testing round, providing insights into the data distribution and variability.

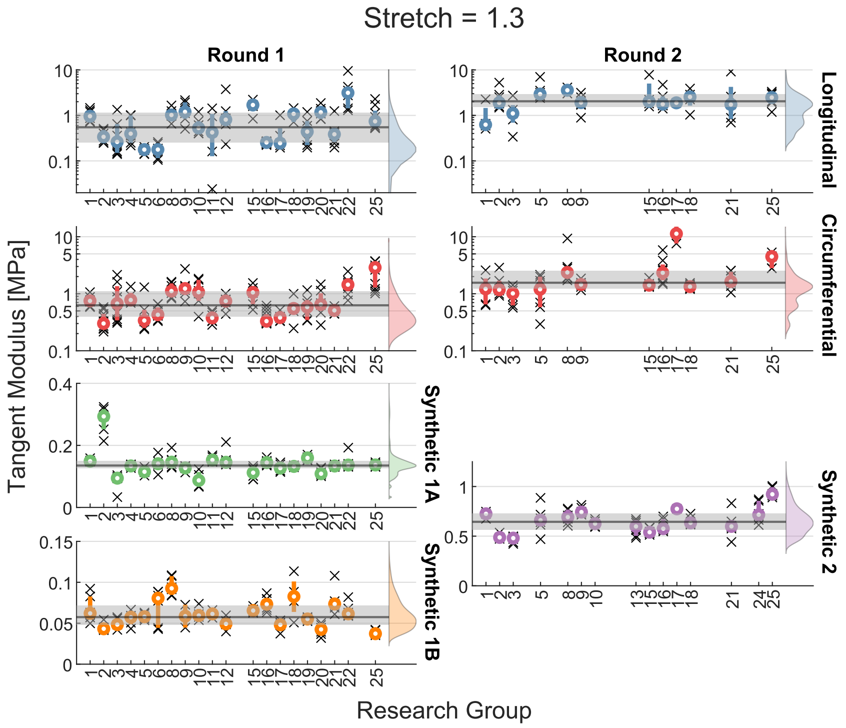


Figure S11: Tangent modulus of the stress-stretch curve at 1.3 stretch per research group, organized by testing round and specimen type. Individual moduli are represented by '×' markers, while the mean for each research group is indicated by a dot and first- to third quantile (Q1-Q3) values are shown as bars. The overall mean and Q1–Q3 values, aggregated across all research groups, are depicted by a line and shaded band, respectively. A density plot is included to represent the distribution of all moduli for each specimen type and testing round, providing insights into the data distribution and variability.

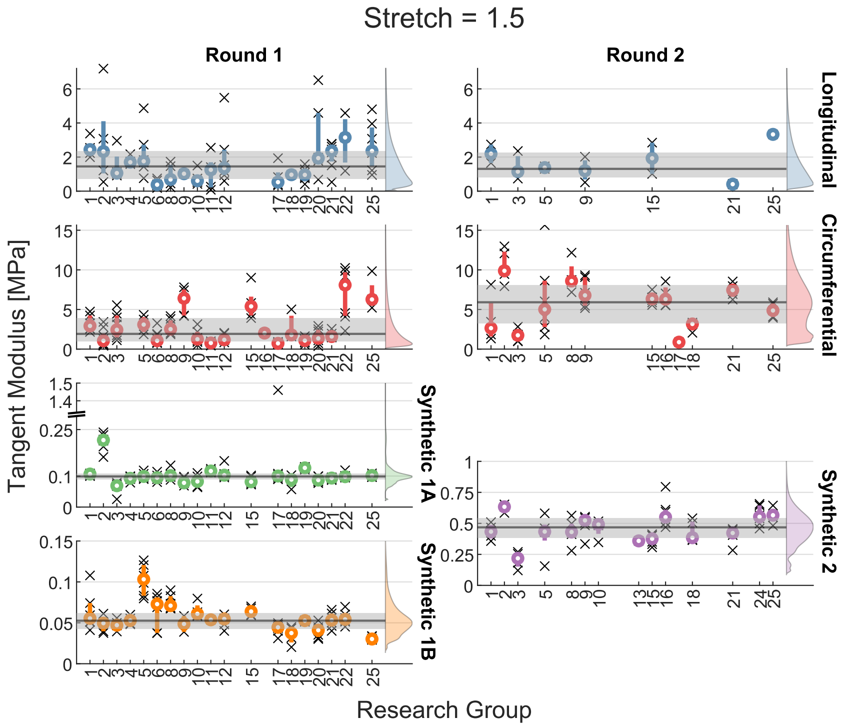


Figure S12: Tangent modulus of the stress-stretch curve at 1.5 stretch per research group, organized by testing round and specimen type. Individual moduli are represented by '×' markers, while the mean for each research group is indicated by a dot and first- to third quantile (Q1-Q3) values are shown as bars. The overall mean and Q1–Q3 values, aggregated across all research groups, are depicted by a line and shaded band, respectively. A density plot is included to represent the distribution of all moduli for each specimen type and testing round, providing insights into the data distribution and variability.

#### Summary of parameters over all groups

Table S1: Summary of parameters by specimen type and round (mean ± SD) (Thickness, Width, Tangent modulus (TM) at a stretch of 1.2 and 1.4, Stress at failure, Stretch at failure and the Numbers of samples tested (N)). From top to bottom first the biological longitudinal samples (L) followed by the biological circumferential samples (C) and the bottom rows represent the synthetic samples (S). Every time first the results from the first round (1) followed by the second round (2).

| | Thickness [mm] | Width [mm] | $TM_{1.2}$ [MPa] | $TM_{1.4}$ [MPa] | Stress* [MPa] | Stretch* [-] | N [-] |
|---|---|---|---|---|---|---|---|
| L1 | 1.8 ± 0.5 | 4.2 ± 1.7 | 0.4 ± 0.6 | 1.4 ± 1.3 | 0.7 ± 0.4 | 1.7 ± 0.3 | 132 |
| L2 | 2.1 ± 0.4 | 4.0 ± 0.2 | 2.0 ± 1.6 | 2.3 ± 1.5 | 0.8 ± 0.3 | 1.4 ± 0.2 | 144 |
| C1 | 2.0 ± 0.5 | 3.9 ± 1.5 | 0.5 ± 0.4 | 1.8 ± 1.9 | 1.3 ± 0.7 | 1.8 ± 0.3 | 117 |
| C2 | 2.4 ± 0.4 | 3.8 ± 0.2 | 2.1 ± 2.6 | 3.7 ± 2.0 | 1.8 ± 0.6 | 1.6 ± 0.2 | 117 |
| S1A | 2.0 ± 0.3 | 3.9 ± 1.9 | 0.2 ± 0.1 | 0.1 ± 0.1 | 0.2 ± 0.1 | 3.4 ± 2.1 | 80 |
| S1B | 2.1 ± 0.2 | 3.8 ± 1.8 | 0.1 ± 0.0 | 0.1 ± 0.0 | 0.2 ± 0.1 | 4.7 ± 3.3 | 88 |
| S2 | 2.0 ± 0.2 | 4.1 ± 0.2 | 0.8 ± 0.2 | 0.5 ± 0.1 | 0.5 ± 0.1 | 1.8 ± 0.3 | 80 |

#### Coefficient of Variation of Tangent Modulus

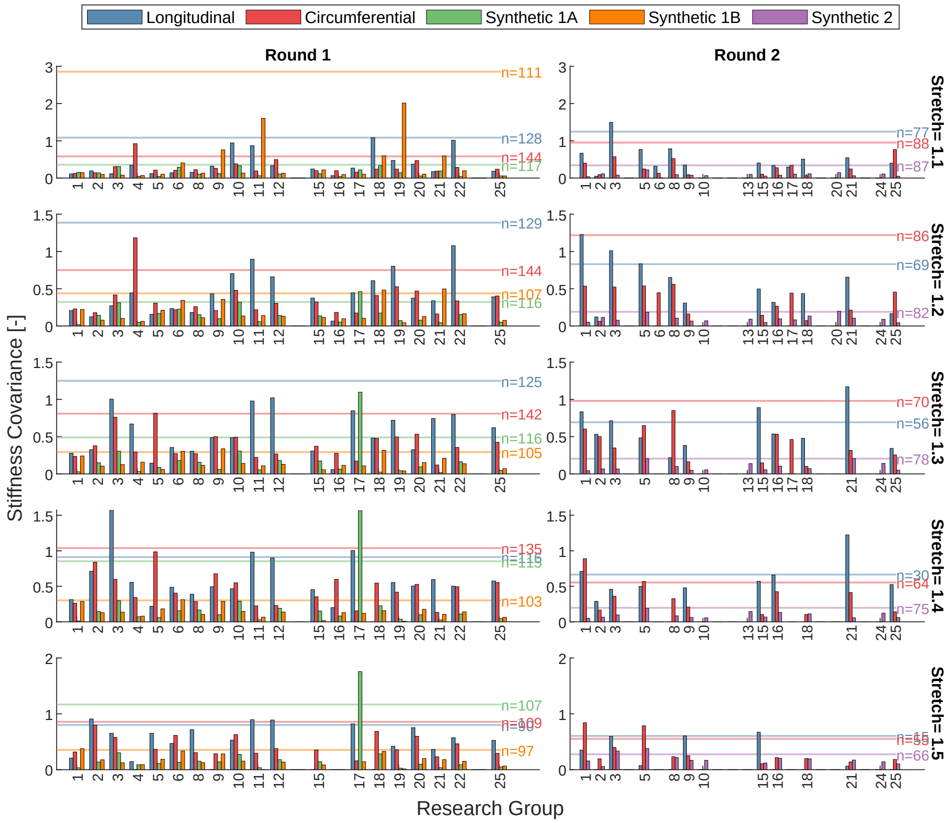


Figure S13: Bar plots showing the coefficient of variation (CV) of the tangent modulus of the stress-stretch curve at 1.1 to 1.5 stretch values with 0.1 steps. CV values are represented for each research group across different specimen types, separated by testing round. The overall CV value, calculated by aggregating measurements across all research groups for each specimen type, is represented by a line. The corresponding sample size is annotated adjacent to the line.
